# Nutrient Microenvironments Reprogram Pigment Epithelium Metabolism and Phenotype

**DOI:** 10.64898/2026.02.11.705448

**Authors:** Rayne R. Lim, Emily Zhao, Daniel T. Hass, Yekai Wang, Mark Eminhizer, Davide Ortolan, Sheldon Niernberger, Aspen Tong, Bree R. Nelson, Marcos Nazario, Vishya Adipudi, Kapil Bharti, James B. Hurley, Jianhai Du, Jennifer R. Chao

## Abstract

Induced pluripotent stem cell-derived retinal pigment epithelium (iPSC RPE) has become a widely used model for studying the mechanisms of age-related macular degeneration (AMD). However, the nutrient composition of currently used RPE culture media is highly variable, posing a major challenge to reproducibility in RPE metabolism and phenotype. We systematically investigate how six distinct nutrient microenvironments shape RPE phenotype, function and metabolism in both iPSC RPE and fetal RPE (fRPE). These included MEMα, DMEM-HG/F12 basal media, physiological human plasma-like medium (HPLM) supplemented with FBS or B27, and X-VIVO 10. Although canonical RPE markers were expressed across all conditions, B27 supplementation and X-VIVO 10 increased RPE cell size, hexagonality, and transepithelial resistance. Culture in HPLM+FBS induced accumulation of lipid droplets and sub-RPE deposits, whereas X-VIVO 10 resulted in the formation of large intracellular vacuoles. B27 supplementation enhanced basal respiration, while X-VIVO 10 increased glycolytic capacity. Amino acid consumption was broadly conserved across media types, including complete depletion of proline in all conditions by 48 hours; however, lipid and nucleotide metabolism varied substantially between conditions. Notably, B27 supplementation in specific media types reversed the net direction of several metabolites, with creatine, serine and taurine shifting from consumption to production, while riboflavin and guanine shifted from production to consumption. These findings establish the nutrient environment as a key determinant of RPE phenotype, function and metabolism. Our work provides a valuable resource for media selection and interpretation of cellular and metabolic phenotypes relevant to RPE disease modeling.

## INTRODUCTION

Age-related macular degeneration (AMD) is a leading cause of visual impairment and blindness in adults over the age of 65 and is projected to affect ∼288 million people worldwide by 2040.(*1*) Preventing degeneration of the retinal pigment epithelium (RPE), which leads to geography atrophy (GA) and central vision loss in AMD patients, remains a major challenge. AMD pathogenesis is complex and multifactorial, and progress has been hindered by lack of ideal animal models. Induced pluripotent stem cell (iPSC)-derived RPE generated from AMD patients and individuals with phenotypically similar monogenic diseases recapitulate several elements of AMD pathology, including sub-RPE deposits resembling basal laminar drusen, dysregulated complement activity, and mitochondrial dysfunction.(*2–8*) Several studies of AMD iPSC and primary RPE models have identified potential disease mechanisms, including mitochondrial injury, susceptibility to oxidative stress, dysregulated lipid metabolism and intracellular complement activation of mTOR signaling pathways.(*7, 9–12*)

While *in vitro* RPE systems show promise for uncovering disease mechanisms and evaluating therapeutic targets, reproducibility and physiologic fidelity remain significant challenges. Contributing factors include variability in iPSC source material, differentiation protocols, and culture conditions.(*13–15*) The nutrient environment, in particular, can exert wide-ranging effects on cellular metabolism and phenotype. Studies in cancer biology have shown that supraphysiologic metabolite concentrations in traditional culture media alter metabolic pathways including glucose utilization, redox state, and gene expression, limiting the ability of cells to mimic in vivo physiology.(*16–18*) Standard media used for iPSC RPE culture have traditionally been optimized for proliferation rather than physiologic relevance. Commonly used supplements such as B27 and N2, originally designed to support neuronal survival, have not been systematically evaluated for their metabolic effects on RPE, although there is evidence that their supplementation reduces mTOR activation in primary RPE.(*19*) More recently, physiologic formulations such as Human Plasma-Like Medium (HPLM) have been shown to rewire cancer cell metabolism to more closely approximate the metabolic profile of primary tumors.(*16, 17*)

Our review of recent human RPE culture studies reveals substantial variability in the nutrient and metabolite composition of commonly used media. Many contain supraphysiologic concentrations of metabolites, including glucose and glutamine, that prevent depletion between media changes but may significantly alter basal RPE metabolism. Our group and others have measured metabolite usage, glycolysis, mitochondrial function, and lipid metabolism across a range of iPSC RPE model systems.(*3, 4, 9, 20*) However, without a consensus on standard media usage and a limited understanding of how medium composition influences RPE metabolism, comparisons across studies and reproducibility remain challenging. Here, we systematically investigate how distinct nutrient environments influence human iPSC and primary fetal RPE (fRPE) morphology, metabolism, and function. By defining how medium composition and supplement choices shape these properties, we aim to improve cross-study interpretability and provide a resource for pathway-specific investigations.

## RESULTS

### Media composition varies widely across RPE culture systems

RPE are cultured under a wide range of nutrient conditions that differ substantially across published iPSC RPE disease modeling studies. To systematically assess this variability, we surveyed reported culture systems and categorized them according to the most frequently used media formulations (**Supp. Table 1**). We observed substantial heterogeneity in nutrient composition, supplement use, and substrate coating matrices. For example, glucose concentrations ranged from physiologic (5 mM) to supraphysiologic (25 mM) levels. Traditional media are often formulated with nonphysiologic nutrient concentrations to prevent depletion between media changes; however, these conditions can markedly alter the metabolic profile of cultured cells.(*16, 17, 21*)

Physiologically relevant cell culture media, such as HPLM (*16*) and Plasmax (*16, 17*) (Ximbio, London, UK), were designed to approximate human plasma metabolite concentrations and more closely recapitulate *in vivo* cellular metabolism and phenotype. The most commonly used traditional media, as well as HPLM with and without B27 supplementation, are shown in **Fig. 1A**, and their reported nutrient compositions are compared in **Supp. Table 2**. MEMα-based media, originally used for human fetal RPE culture (*22*), contain relatively high levels of growth factors (e.g., insulin, transferrin, and progesterone) and amino acids (e.g., cysteine, aspartate, arginine, and proline). DMEM/F12 supplemented with 2% B27 provides higher levels of essential fatty acids (e.g., linoleic and linolenic acids), amino acids (e.g., leucine, lysine, and serine) and antioxidants (e.g., catalase and superoxide dismutase). The X-VIVO 10 formulation is proprietary; however, available information indicates that glucose and glutamine concentrations are the highest among the media examined. HPLM, which has not previously been applied to RPE culture, contains physiologic plasma concentrations of glucose, amino acids, and additional metabolites readily utilized by RPE such as lactate and succinate, but lacks exogenous growth factors typically included in RPE media. Given this variability, we next examined how differences in nutrient and growth factor composition influence RPE phenotype in culture.

**Figure 1.**
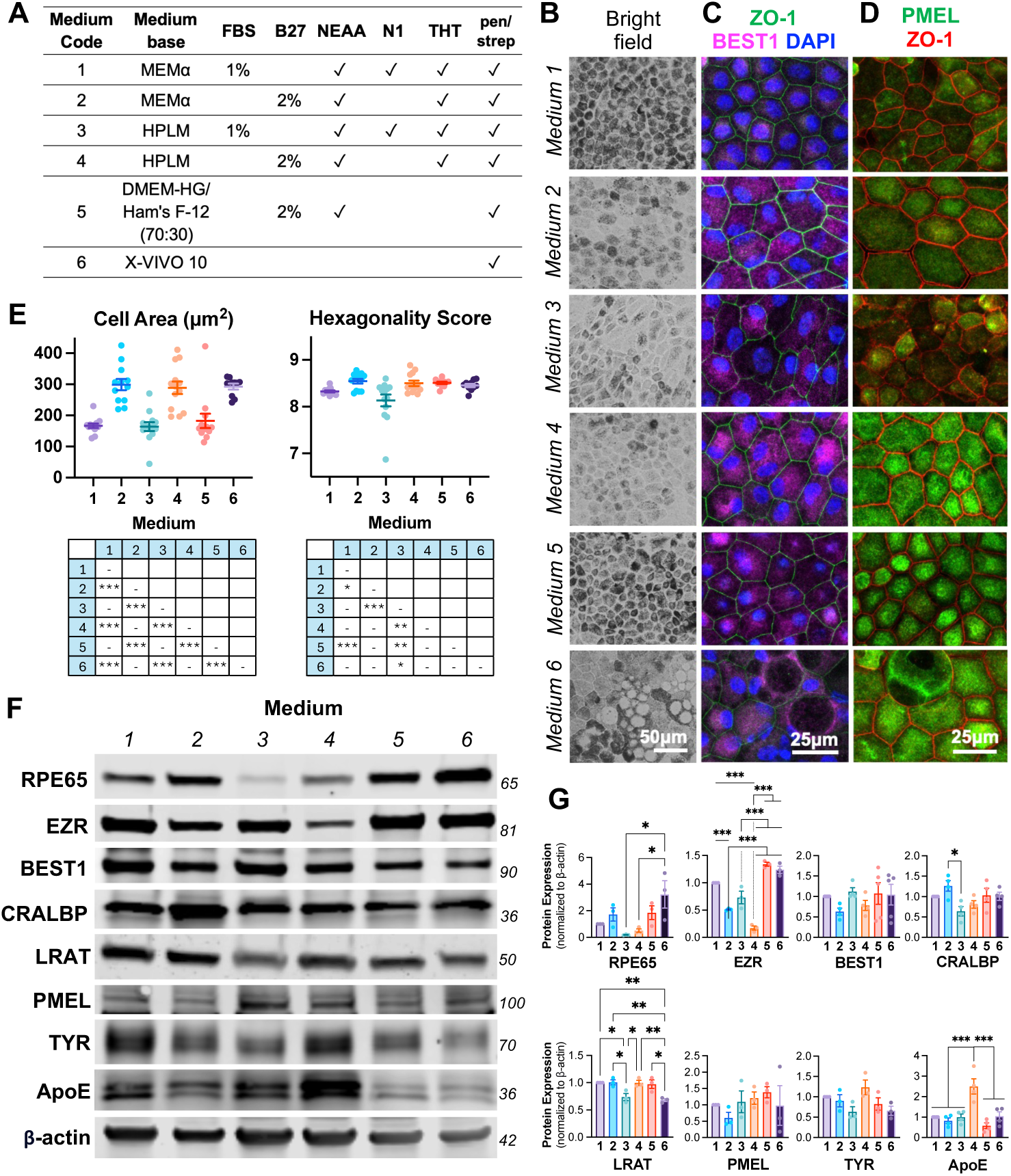
**iPSC RPE morphology and marker expression is altered by media composition.**(**A**) Formulation of Media 1 – 6 used in this study. (**B**) Representative brightfield images of iPSC RPE cultured for up to 8 weeks on Matrigel coated culture dishes. (**C,D**) iPSC RPE cultured on chambers slides were fixed and immunostained for ZO-1, BEST1, and PMEL. (**E**) RPE cell area and hexagonality were calculated using REShAPE analysis. (**F,G**) Immunoblotting and quantification of key RPE markers in RPE lysate after 8 weeks culture in Media 1 – 6. Mean ± SEM. *, p<0.05; **, p<0.01; ***, p<0.001.

### Nutrient composition influences RPE phenotype and morphology

iPSC RPE and fRPE were cultured on Matrigel-coated filter inserts in six distinct media formulations (**Fig. 1A, Supp. Fig. 1A**) for 8-10 weeks. iPSC RPE exhibited significant differences in pigmentation and polygonal morphology across conditions (**Fig. 1B**). ZO-1 immunostaining, indicating the presence of tight junctions, was detected in all conditions (**Fig. 1C**). Quantitative analysis of cell surface area and hexagonality with ZO-1 and F-actin staining using REShAPE software showed that iPSC RPE in media 2, 4 and 6 were approximately 60% larger than those in media 1, 3, and 5. Media 2 and 4 contained 2% B27 supplementation, whereas media 1 and 3 did not; medium 5 also contained B27 but differed in base medium composition. Notably, cells cultured in HPLM supplemented with FBS (medium 3) exhibited the lowest degree of hexagonality (**Fig. 1E, Supp. Fig. 1H**).

Expression of key RPE markers, including BEST1, PMEL and Tyr, was maintained across all conditions (**Fig. 1F-G**). BEST1 immunostaining revealed similar distribution patterns across medium types (**Fig. 1C**), whereas PMEL staining appeared more uniform and intense in B27-supplemented media (media 2, 4, and 5) and in X-VIVO 10 (medium 6) (**Fig. 1D**). In contrast, RPE65 expression was significantly reduced in HPLM, while EZR levels were elevated in DMEM-HG and X-VIVO 10. LRAT expression was lower in HPLM supplemented with FBS and in X-VIVO 10 (**Fig. 1F-G**). Comparable, medium-dependent differences were also observed in fRPE (**Supp. Fig. 1B-D**).

### Nutrient composition influences RPE polarity and barrier function

Transepithelial resistance (TER) increased with RPE maturation, exceeding 200 Ω⋅cm^2^ by four weeks of culture in all media conditions except medium 3 (**Fig. 2A**). The magnitude of TER varied considerably across conditions, ranging from <200 Ω⋅cm^2^ to nearly 800 Ω⋅cm^2^, with X-VIVO among the highest. Notably, replacement of FBS with B27 significantly enhanced TER, consistent with increased tight junction formation. In addition, RPE cultured in media 3 and 5 consistently exhibited higher basal VEGF secretion on both polyester (PET) and mixed cellulose ester filter inserts (**Supp. Fig. 2A-B**). When assessed in RPE cultured on flat-bottom plates, VEGF expression was more variable between iPSC RPE (**Fig. 2B**) and fRPE (**Supp. Fig. 2C**) but overall showed similar media-dependent trends.

**Figure 2.**
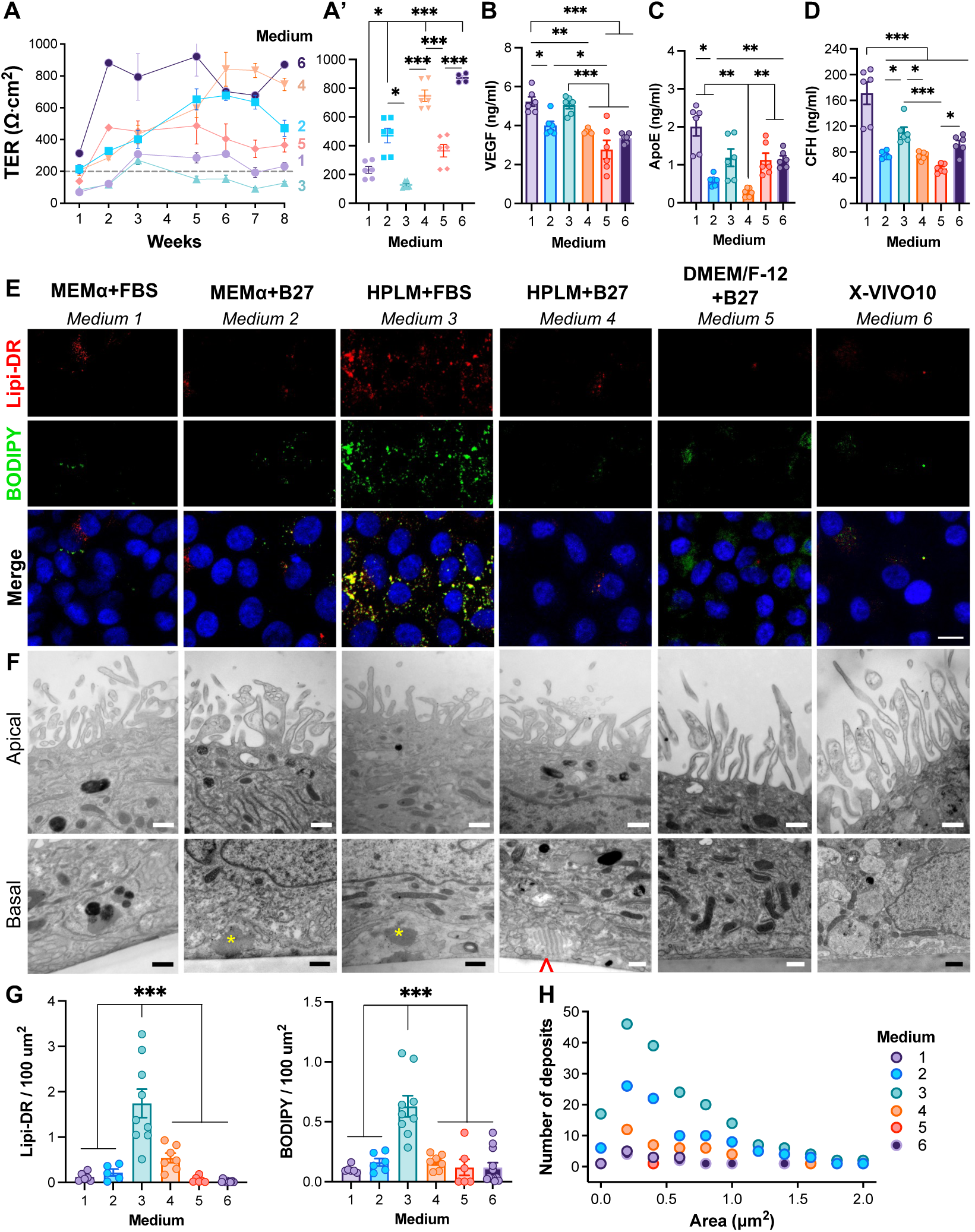
Nutrient composition influences RPE polarity and sub-RPE deposition. (**A**) TER of iPSC RPE over time on 0.33cm^2^ PET Transwell filters, and (**A’**) at 8 weeks. (**B-D**) ELISA of VEGF, ApoE and CFH in 24h conditional media from RPE cultured on flat bottom dishes. Mean ± SEM. *, p<0.05; **, p<0.01; ***, p<0.001. (**E**) Live cell stain with Lipi-DeepRed (Lipi-DR) and BODIPY-493/503 to identify lipid droplets in the 8 weeks iPSC RPE. Scale bar, 10μm. (**F**) Representative TEM images of apical microvilli and basal infoldings with sub-RPE deposits in 8 weeks iPSC RPE on Transwell filters. *, amorphous deposits; ^, banded collagen. Scale bar, 500nm. (**G**) Quantification of lipid droplets per 100μm^2^ area in Lipi-DR and BODIPY images. (**H**) Sub-RPE deposits were identified and quantified in n=3 panoramic sections of the TEM images.

Because RPE cell size and density varied across cultured conditions, both nuclei counts and total protein measurements are provided for iPSC RPE cultured on filter inserts, flat-bottom chambers, and Seahorse plates across all media types to aid interpretation (**Supp. Fig. 2D**). With the exception of RPE maintained in X-VIVO 10, nuclei counts were generally consistent between RPE cultured on filter inserts and flat-bottom dishes. However, nuclei count did not reliably correspond to total intracellular protein, as higher nuclei numbers did not consistently reflect greater protein content, likely due to differences in cell size and/or intracellular protein density. Consequently, normalization to either total protein (**Supp. Fig. 2E’**) or nuclei count (**Supp. Fig. 2E”**) produced divergent results. To avoid introducing bias through normalization, data in this study are presented without normalization unless otherwise indicated.

### Nutrient composition alters lipid accumulation, complement activity, and autophagy in RPE

Modeling AMD *in vitro* is a valuable approach for investigating disease-relevant mechanisms. We next examined whether alterations in the nutrient environment influence AMD-associated features such as lipid accumulation and sub-RPE deposit formation. Neutral lipids were visualized using the fluorescent live-cell sensors Lipi-DR and BODIPY. RPE maintained in HPLM supplemented with FBS exhibited the highest levels of lipid accumulation (**Fig. 2E**), which were significantly higher compared with all other media conditions (**Fig. 2G**). By electron microscopy, RPE cultured for 8 weeks in all media conditions developed apical processes and basal infoldings (**Fig. 2F**). Basal RPE deposits, including amorphous material or banded collagen, were most prevalent in iPSC RPE maintained in HPLM supplemented with FBS, followed by MEMα supplemented with B27 (**Fig. 2H**). Notably, apical microvilli in B27-supplemented media (media 2 and 4) contained more electron lucent vesicles than RPE cultured in the corresponding base media supplemented with FBS.

Using ApoE as a proxy for lipoprotein production, both iPSC RPE and fRPE cultured in MEMα+B27 and HPLM+B27 secreted less ApoE than their FBS-supplemented counterparts (**Fig. 2C, Supp. Fig. 2C**). Complement-related changes were likewise medium dependent. Complement factor H (CFH) production secretion was highest in medium 1 and lowest in B27-containing media (media 2, 4, and 5) (**Fig. 2D, Supp. Fig. 2C**). Intracellular CFH was also examined by immunostaining in iPSC RPE under all conditions (**Supp. Fig. 3A**). Clusterin (CLU) was reduced in iPSC RPE cultured in HPLM+B27 (**Supp Fig. 3A**), although this was not consistent in fRPE, which showed increased CLU with B27 supplementation (**Supp. Fig. 1E**). CD46, an inhibitor of the alternative pathway of the complement system, showed a shift in localization from diffuse cytosolic staining under B27-supplemented conditions to increased plasma membrane association under high-glucose conditions (**Supp. Figs. 1E and 3A**).

Markers of autophagy, a pathway whose dysregulation is implicated in RPE degeneration in AMD, also varied by medium type. Expression of Beclin1, an autophagy-related protein, was reduced in iPSC RPE cultured in MEMα+B27 and HPLM+B27 media (**Supp. Fig. 1F and 3B**). B27 supplementation reduced LAMP1 (lysosome-associated membrane protein) expression in MEMα-based media but increased LAMP1 levels in HPLM in both iPSC RPE and fRPE (**Supp. Figs. 1G and 3C**). Expression of both Beclin1 and LAMP1 was generally lower in RPE cultured in X-VIVO 10. Together, these findings suggest that AMD-relevant pathways, including lipid homeostasis, complement regulation, and autophagy are influenced by the nutrient environment.

### Nutrient composition affects oxygen consumption and glycolytic capacity

Substrate availability directly influences mitochondrial tricarboxylic acid (TCA) cycle activity. After iPSC RPE and fRPE were plated in XFe96 well plates and cultured in their respective media for four weeks, they were transitioned to Seahorse XF DMEM for 1 hour prior to assessment of mitochondrial and glycolytic function using the Seahorse XF analyzer. B27 supplementation increased basal respiration and ATP-linked respiration in iPSC RPE but did not significantly affect maximal respiration (**Fig. 3A**). In both iPSC RPE and fRPE (**Fig. 3A, Supp. Fig. 4A**), cells cultured in medium 3 exhibited the lowest respiratory activity as well as spare capacity, possibly reflecting its less mature RPE-like phenotype. As noted earlier, because total protein content did not consistently correlate with nuclei counts (**Supp. Fig. 4C-D**), oxygen consumption rate (OCR) values normalized to each parameter are shown for comparison (**Supp. Fig. 4E-F**). Glycolysis and glycolytic capacity did not differ significantly among media types 1-5, whereas X-VIVO 10 increased both basal glycolysis and glycolytic capacity in iPSC RPE (**Fig. 3B**), indicating a selective effect of this medium on glycolytic metabolism. B27 supplementation significantly increased glycolytic reserve and non-glycolytic acidification in media 2 and 4 (**Fig. 3B, Supp Fig. 4B**). Non-glycolytic acidification was highest in iPSC RPE and fRPE cultured in DMEM/F-12, suggesting reliance on other fuel sources in the nutrient rich medium.

**Figure 3.**
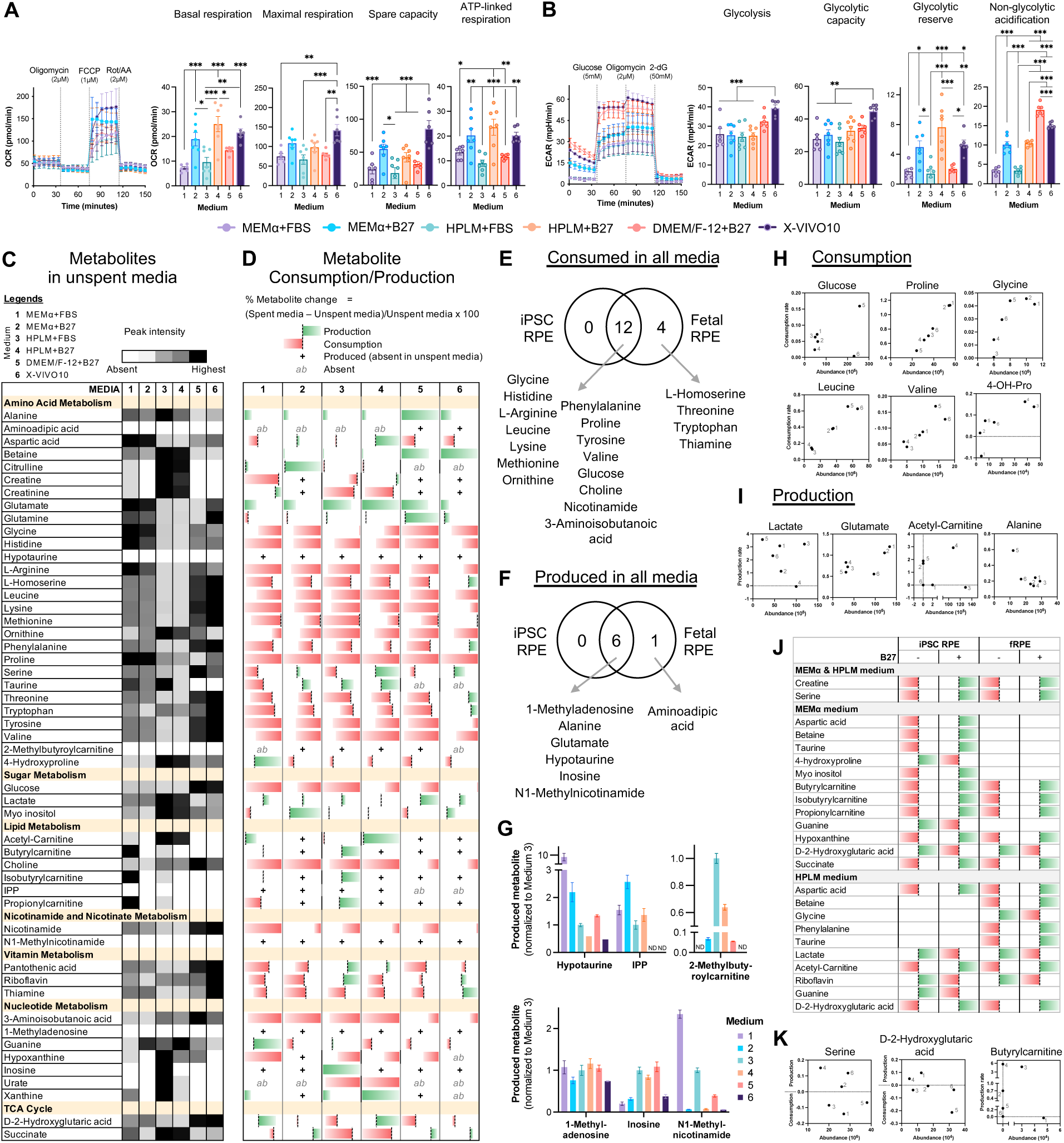
Respiration, glycolysis and extracellular metabolite consumption/production in iPSC RPE. (**A,B**) iPSC RPE were seeded and maintained on Matrigel coated XFe96 well plates in respective Media 1 – 6 for four weeks. Prior to (**A**) mitochondrial stress test and (**B**) glycolysis stress test, cells were equilibrated in XF DMEM media (pH 7.4) in CO_2_-null incubator for 1 hour. Mean ± SEM. *, p<0.05; **, p<0.01; ***, p<0.001. (**C-K**) Targeted metabolomics on 8-week-old iPSC RPE was performed via LCMS. (**C**) Extracellular metabolites in unspent media were plotted as a heat map, with degrees of grey depicting lowest to highest intensity. (**D**) To calculate metabolite changes in 48h spent media from iPSC RPE, results are depicted on a sliding scale where % metabolite = (Spent media – Unspent media)/Unspent media x 100. Green, production; Red, consumption; **+**, new production; *ab*, absent. Venn diagram showing common and unique metabolites (**E**) consumed or (**F**) produced across all media between iPSC RPE and fRPE. (**G**) Newly produced metabolites that were initially absent in unspent media normalized to Medium 3. (**H,I**) Consumption rate and production rate of select metabolites plotted against abundance in unspent media. Media number noted in graph in grey. (**J,K**) Comparison of Medium 1 vs 2 and Medium 2 vs 3, to show changes in usage of metabolites when FBS is replaced with B27.

**Figure 4.**
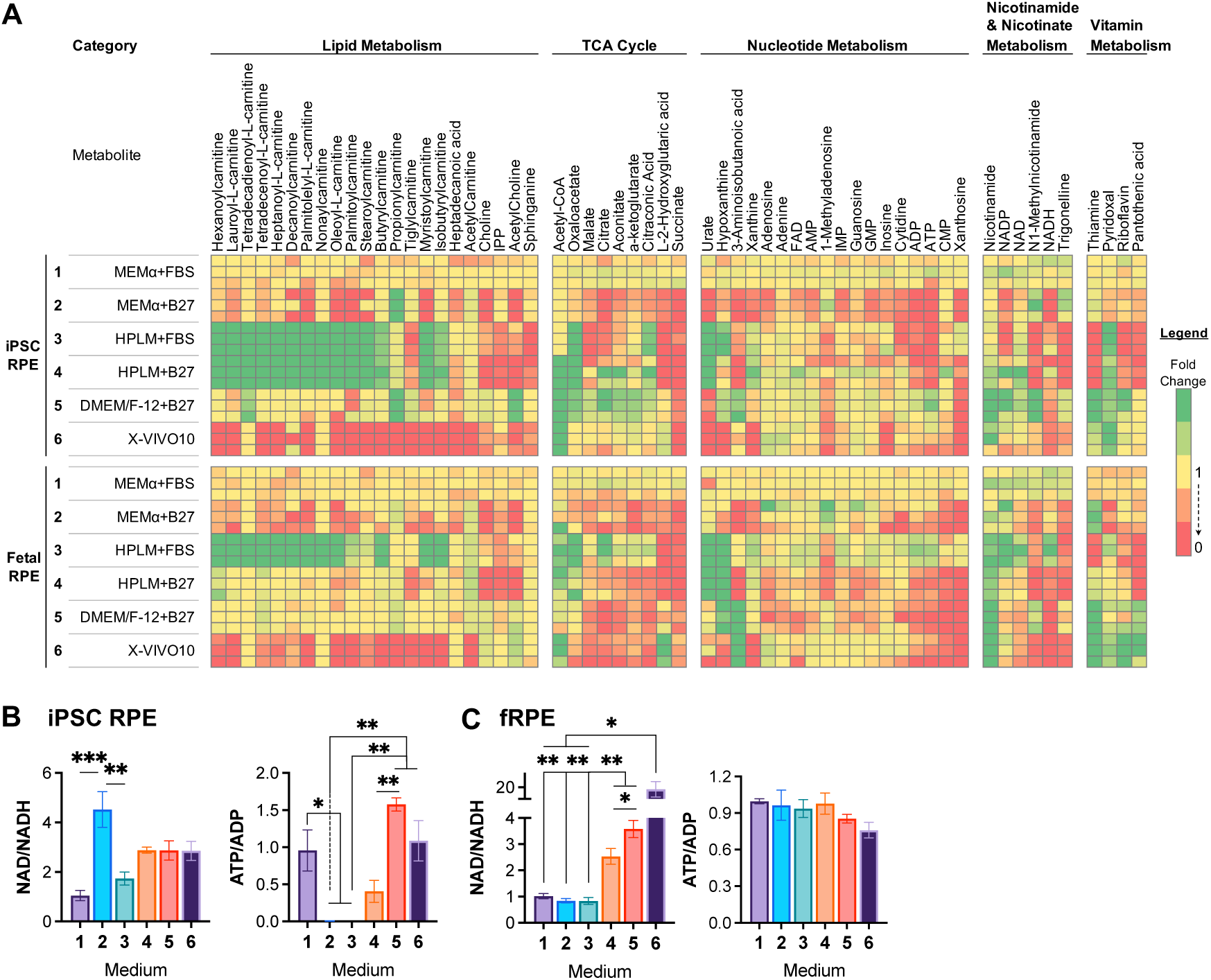
Heatmap of intracellular metabolites involved in lipid, nucleotide, NAD, vitamin metabolism and TCA cycle in iPSC RPE and fRPE. (**A**) 114 detected intracellular metabolites were categorized by major metabolic pathways, sorted left to right from highest to lowest abundance, normalized to Medium 1 and depicted as a heatmap. Yellow, fold change (FC)=1; Red, FC<1; Green, FC>1. Ratio of NAD/NADH and ATP/ADP in lysate of (**B**,**C**) iPSC RPE and (**D**,**E**) fRPE maintain in Media 1 – 6 at 8 weeks. Mean ± SEM. *, p<0.05; **, p<0.01; ***, p<0.001.

### RPE metabolism is influenced by culture media composition

Of the 378 targeted metabolites, we detected 51 extracellular and 114 intracellular metabolites, with 29 overlapping reported media components (**Fig. 3C**). Multivariate analysis using partial least squares discriminant analysis showed clear separation among media types in both cell lysates and conditioned media (**Supp. Fig. 5A**), indicating that the RPE metabolome is highly dependent on culture medium composition (**Supp. Fig. 5B**).

To assess metabolite production and consumption, values measured in unspent media were subtracted from those in 48-hour spent media and mapped on a color scale, where green denotes production and red denotes consumption. Extracellular metabolic profiles were generally consistent between iPSC RPE (**Fig. 3D**) and fRPE (**Supp. Fig. 5C**). Both consumed amino acids including leucine, lysine, valine, proline and ornithine across all media conditions (**Fig. 3E**), while producing alanine, glutamate and hypotaurine (**Fig. 3F**). Several metabolites, including proline, leucine, valine and glycine, were consumed in an availability-dependent manner, although glucose consumption differed in media 5 and 6 despite similarly high starting concentrations (**Fig. 3H**). Glutamate production rate was highest in media 1 and 2, which also had the highest unspent media abundance, whereas alanine production increased under low-abundance conditions. B27-containing media (2, 4, and 5) induced net acetyl-carnitine production, whereas HPLM+FBS (medium 3) resulted in net consumption (**Fig. 3I**).

Several metabolites absent from unspent media were highly produced by iPSC RPE, including N1-methylnicotinamide and hypotaurine, which were most abundant in medium 1, and 2-methylbutyroylcarnitine, which increased nearly 20-fold in medium 3 compared with medium 2 (**Fig. 3G**). Media-specific differences were also observed. Urate, present only in HPLM-based media, was highly consumed, whereas hypoxanthine was consumed when supplied but produced when absent. 4-hydroxyproline, a by-product from collagen turnover (*23, 24*), was produced exclusively in MEMα+FBS (**Fig. 3D, 3H**). Replacement of FBS with B27 in either MEMα- or HPLM-based media inverted the net usage of numerous metabolites (**Fig. 3J**). For example, creatine and serine were consumed in MEMα and HPLM supplemented with FBS but became net produced under B27-supplemented conditions. Similar reversals were observed for 4-hydroxyproline, guanine and D-2-hydroxyglutaric acid, which shifted from net production to consumption in MEMα supplemented with B27, compared to MEMα supplemented with FBS (**Fig. 3K**). Although iPSC RPE and fRPE exhibited broadly similar metabolite consumption and production patterns across media conditions, a few differences were observed (**Supp. Fig. 5D)**. For instance, betaine (medium 1) and glycine (medium 3) were consumed by iPSC RPE but produced by fRPE, whereas glutamine in media 3 and 4 were produced by iPSC RPE but consumed by fRPE.

Intracellular metabolites were grouped by pathway, ranked by relative abundance compared with medium 1 (highest to lowest, from left to right), and visualized by heatmap (**Fig. 4 and 5**). In lipid metabolism, both iPSC RPE and fRPE cultured in X-VIVO 10 had the lowest levels of acylcarnitines, whereas those cultured in HPLM+FBS had the highest. Replacement of FBS with B27 preserved acylcarnitine levels in iPSC RPE cultured in medium 4, while both iPSC RPE and fRPE exhibited reduced intracellular choline, acetylcholine and isopentenyl pyrophosphate levels (**Fig. 4A**). Medium 2 showed the lowest accumulation TCA cycle and nucleotide intermediates in both iPSC RPE and fRPE. Interestingly, redox state and its coupling with cellular energy status differed between cell types. iPSC RPE cultured in medium 2 had an increased NAD^+^/NADH ratio and a decreased ATP/ADP ratio (**Fig. 4B**), whereas fRPE showed increased in NAD^+^/NADH in media 4-6, without a corresponding change in ATP/ADP ratio (**Fig. 4C**).

**Figure 5.**
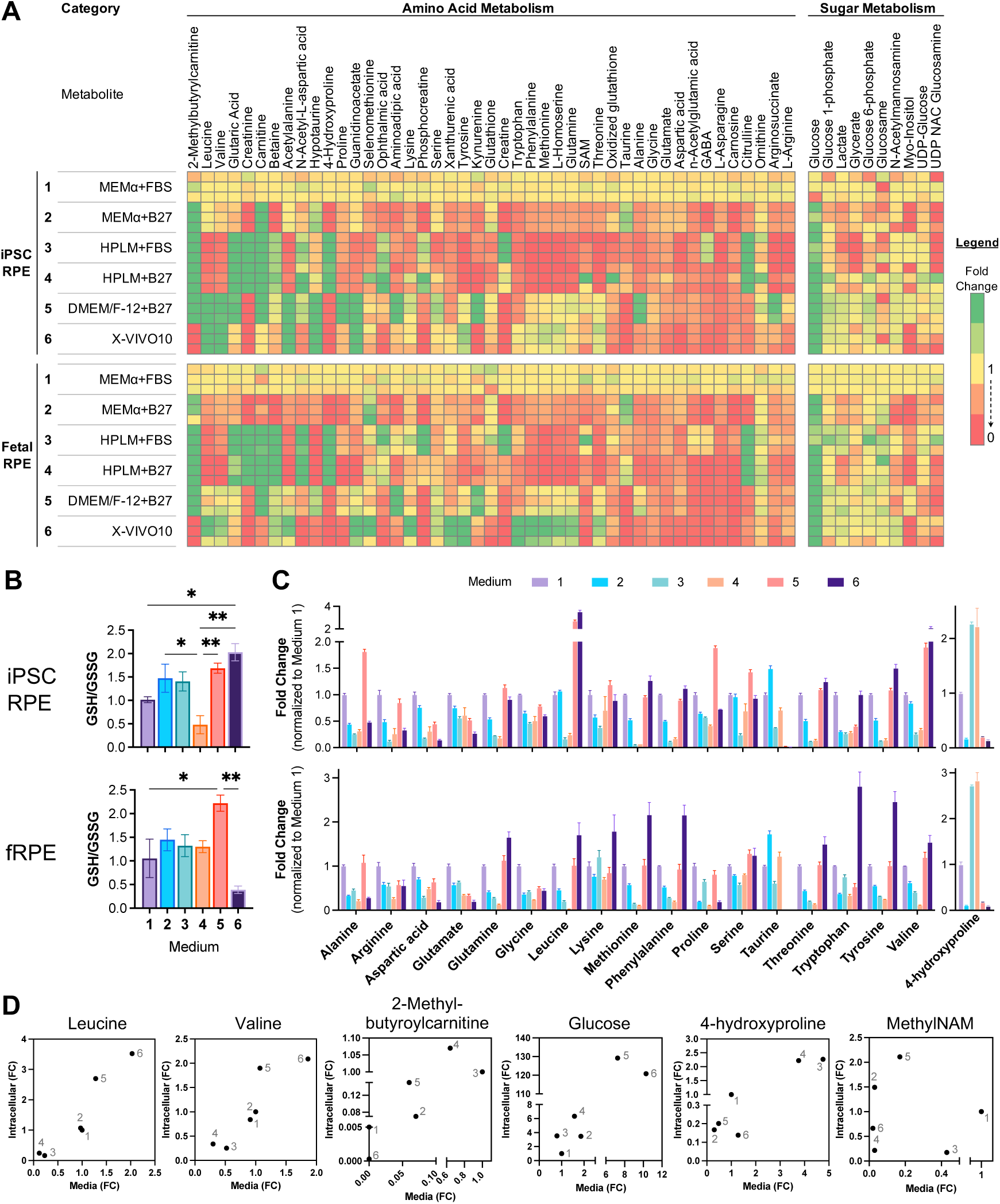
Intracellular metabolites of amino acid and sugar metabolism. (**A**) Heatmap of metabolites are analyzed as described in Figure 5. (**B**) Ratio of GSH/GSSG in iPSC RPE and fRPE. (**C**) Histogram of amino acid abundance normalized to medium 1. (**D**) Select intracellular metabolites were plotted against extracellular abundance in 48h spent media. All data normalized to medium 1, except for 2-methyl-butyroylcarnitine, which was normalized to medium 3. Mean ± SEM. *, p<0.05; **, p<0.01.

Amino acid utilization varied with medium composition (**Fig. 5A, 5C**). In both iPSC RPE and fRPE, intracellular amino acid levels were lowest in media 3 and 4 and highest in media 5 and 6, broadly reflecting starting media concentrations. Likewise, intracellular glucose levels were highest in RPE cultured in media 5 and 6, which contained the highest starting glucose concentrations. In both RPE cell types, intracellular leucine and valine abundance closely mirrored their abundance in 48h spent media (**Fig. 5D**). Both branched-chain amino acids (BCAA) were low in media 1-4 but markedly higher in media 5 and 6. BCAA-derived acylcarnitines – 2-methylbutyrolcarnitine, isobutyrylcarnitine (**Fig. 4A**) and propionylcarnitine (**Fig. 4A**) – were lowest in medium 6, and highest in media 3 and 4, suggesting that BCAA utilization is reduced under high extracellular glucose or amino acid conditions. Intracellular 4-hydroxyproline abundance was highest in HPLM-based media 3 and 4 (**Fig. 5C**), which also had the highest starting levels of the substrate (**Fig. 3C**). Interestingly, 4-hydroxyproline was only net produced in medium 1, suggesting that the nutrient environment can also alter extracellular processes, including extracellular matrix turnover in *in vitro* RPE.

To directly assess the metabolic effects of media supplementation, intracellular metabolites were compared between FBS- and B27-supplemented conditions. In both MEMα-and HPLM-based media, B27 supplementation consistently enriched pathways associated with BCAA degradation and the Warburg effect (e.g., enhanced glycolysis and lactate production) (**Supp. Fig. 6B**). These findings suggest that B27 supplementation alters amino acid catabolism and promotes a more glycolytic phenotype in the RPE, independent of the basal medium composition.

Finally, markers of oxidative stress also varied by medium composition. The GSH/GSSG ratio was largely comparable across conditions in iPSC RPE and fRPE, although the ratio was significantly reduced in medium 4 relative to medium 1 in iPSC RPE and in medium 6 in fRPE, indicating increased oxidative stress (**Fig. 5B**).

## DISCUSSION

The RPE is highly sensitive to its nutrient environment, which influences RPE cell morphology, disease-relevant phenotype, and metabolism. Here, we systematically compared six commonly used culture media to determine how nutrient composition and supplement choice impact RPE phenotype and metabolism. Although canonical RPE markers were maintained across all conditions, cell morphology, barrier function, and AMD-relevant features such as lipid accumulation, complement factor secretion, and autophagy markers varied widely. Metabolomic profiling further showed that amino acid, lipid, nucleotide, and redox metabolism are strongly influenced by substrate availability and supplementation. Together, these findings highlight the impact of the nutrient microenvironment on RPE phenotype and metabolism (summarized in **Fig. 6**) and underscore medium composition as a critical variable in the design and interpretation of in vitro RPE disease models.

**Figure 6.**
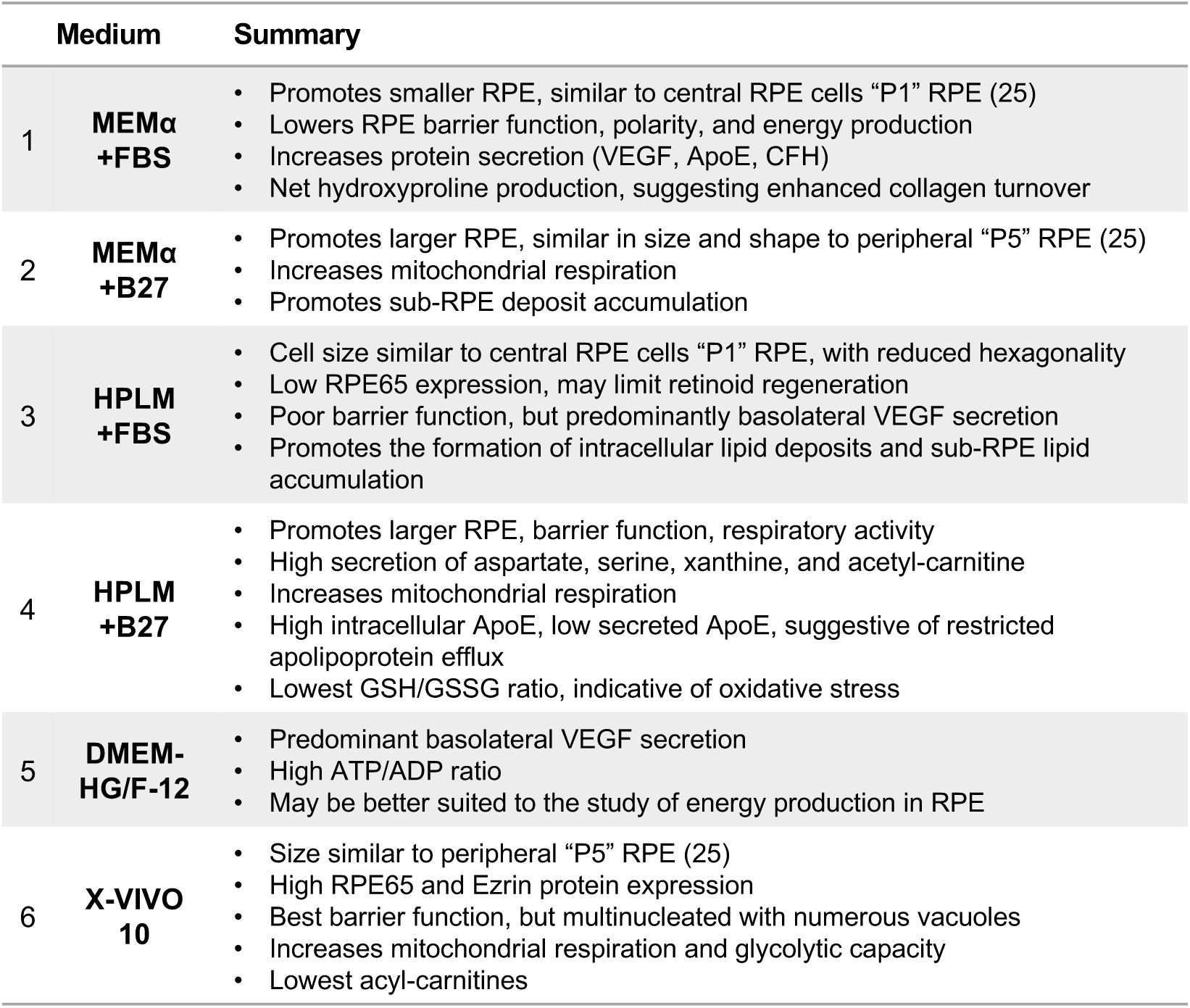
**Summary of RPE Phenotype and Metabolism Across Media Conditions (1 – 6).**

Medium 1 (MEMɑ + 1% FBS) has long been used as a standard formulation for RPE culture (**Supp. Table 1**). RPE maintained in this medium exhibit smaller cell size, similar to central “P1 RPE”, and display reduced hexagonality relative to cells cultured in the same base medium supplemented with B27.(25) Tight junction formation, epithelial polarity, and mitochondrial energy production are generally lower in this medium than in other media types. In contrast, RPE cultured in this medium show increased secretion of CFH, ApoE, and VEGF, although it remains unclear whether this reflects enhanced synthesis or secretion. 4-hydroxyproline release is also elevated, suggesting greater collagen turnover, and this is the only condition that demonstrated net hydroxyproline production. Collectively, the most prominent characteristics of this medium are that it promotes elevated protein secretion and extracellular matrix remodeling by RPE. Medium 2 (MEMɑ + B27) differs from medium type 1 by the substitution of FBS with B27 supplementation. This modification resulted in increased RPE cell size, yielding cells that resemble peripheral “P5 RPE”.(25) The presence of B27 increases mitochondrial respiration in iPSC RPE and promotes the accumulation of sub-RPE deposits, making it useful for modeling aspects of both normal RPE physiology and lipid-related disease processes such as AMD.

Medium 3 (HPLM + 1% FBS) represents the most physiologic formulation, originally developed to approximate human plasma composition in cancer studies(16, 17) and applied here for the first time to RPE culture. Compared to other media, HPLM provides a more physiologic, nutrient rich spectrum, which may contribute to enhanced lipid synthesis and utilization, consistent with the metabolic flexibility of RPE observed previously.(26) When supplemented with serum, HPLM closely approximates the physiologic basal RPE environment immediately following feeding, although the in vivo composition of the subretinal/apical environment remains incompletely defined. Cells cultured in this medium are similar in size to central “P1 RPE”,(25) although with reduced hexagonality, a feature we observed to be common to FBS-containing conditions. Barrier function is relatively poor, possibly reflecting the smaller cell size,(27) and both respiratory activity and glycolytic reserve were lowest in this medium. RPE maintained in this medium exhibit elevated secretion of CFH and ApoE, along with increased release of short-chain acylcarnitines (propionyl-, butyryl-, isobutyryl-, and 2-methylbutyrylcarnitine). Acylcarnitines are in near equilibrium with acyl-CoAs,(28) and the corresponding CoAs are intermediates in isoleucine, valine, and odd-chain fatty acid degradation pathways. Cytosolic lipid accumulation and sub-RPE deposition are significantly elevated, which when taken together with the accumulation of metabolic intermediates associated with amino acid and fatty acid catabolism imply an impairment in fatty acid utilization. The most prominent feature of this medium is its induction of intracellular lipid accumulation and sub-RPE deposit formation, and more than in medium 2, may provide a robust model for studying AMD-related RPE phenotypes. Notably, RPE65 expression is suppressed in this medium, suggesting the most limited retinoid regeneration among media types tested.

Medium 4 (HPLM + B27) substitutes the B27 supplement for FBS. As observed with MEMɑ, this exchange increases iPSC RPE size, enhances barrier integrity, and elevates respiratory activity. Intracellular ApoE levels are highest under this condition, although this increase does not correspond to elevated ApoE secretion. Intermediates of lipid metabolism were retained in the cytosol; however, lipid accumulation was reduced compared to its FBS supplemented counterpart, suggesting that B27 alters fatty acid storage in these conditions. There are few parameters in which this medium is clearly superior or inferior to other types, although some features of RPE are distinctive, including the uniquely high secretion of aspartate, serine, xanthine, and acetyl-carnitine. The functional significance of this however is unclear. Notably GSH/GSSG is lowest in iPSC RPE maintained in this medium, suggesting increased oxidative stress. Overall, HPLM+B27 can be considered an intermediate medium for the parameters we investigated.

Medium type 5 (DMEM/F12 with high glucose) supports RPE with a size comparable to central “P1 RPE”, (25) although with reduced hexagonality. This medium is distinctive in promoting predominantly basal VEGF release, suggesting that its components may contribute to establishing or maintaining normal RPE polarity. DMEM/F12 contains elevated concentrations of glucose (20.5 mM), amino acids (including branched chain amino acids, tyrosine, serine), and choline. Serine supports metabolic flexibility, antioxidant defense, and lipid balance critical to RPE homeostasis(26, 29) and choline is essential for lipid metabolism in RPE, supporting phospholipid synthesis, membrane maintenance, and metabolic crosstalk with photoreceptors. (30) Both serine and choline are integral to one-carbon metabolism, influencing DNA methylation and epigenetic regulation.(31, 32) Production of glutamine and alanine, both important sources of nitrogen for the retina, was increased, suggesting enhanced nitrogen metabolism.(33, 34) RPE cells cultured in this media have the highest ATP/ADP ratio. Overall, DMEM/F12 + high glucose may be best suited for studies probing metabolic regulation and energy production in RPE.

Medium 6 (X-VIVO 10) is the most nutrient-enriched, providing the highest concentrations of amino acids, in addition to high glucose. It also contains pantothenic acid, a precursor for CoA in lipid metabolism, and such abundant nutrients may predispose the RPE to lipid metabolism perturbations. RPE cultured in this medium resemble peripheral “P5 RPE” in size and shape.(25) They exhibit the highest levels of RPE65 among all media types tested, while LRAT levels are lower relative to most other medium conditions, leading us to suspect the greatest retinoid regeneration capacity in this medium. X-VIVO 10 also supports relatively high secretion of CFH, ApoE, and VEGF, without compromised TER, in contrast to medium 1. Despite these potentially favorable features, several atypical morphological features were observed, including multinucleated cells and the accumulation of numerous electron-lucent intracellular vacuoles that appear to contain organelle remnants and multilaminated membranes like autophagic vesicles (**Supp. Fig. 7C**). However, immunostaining with various markers did not further reveal the identity of the vacuoles (**Supp. Fig. 7D**). Melanosomes were noted earlier in the maturation process compared to other media types, consistent with previous reports of pronounced pigmentation in iPSC RPE differentiated and maintained in X-VIVO 10.(35) Overall, this medium may be best suited for studying the RPE component of retinoid cycling, while warranting caution regarding nutrient-associated morphological changes.

B27 is a chemically defined supplement (**Supp. Table 2**.) originally formulated in the 1990s to support neuronal survival and differentiation under serum-free conditions.(36) It improved upon earlier supplements, such as N2, which lacked key nutrients and antioxidants to sustain neuronal viability.(37) B27 was subsequently adopted for RPE culture as a serum-free alternative to reduce experimental variability, eliminate undefined survival factors in FBS, and establish xeno-free culture conditions suitable for clinical-grade manufacturing.(38, 39) In our studies, B27 supplementation produced multiple effects on RPE phenotype, including increased cell size, enhanced hexagonality, and improved tight junction formation as measured by TER, all consistent with a more differentiated epithelial state. In contrast, the undefined growth factors and mitogenic signals present in FBS may promote a more mesenchymal or de-differentiated state. These findings align with prior reports showing that B27 substitution reinforces enhanced epithelial phenotype in RPE.(39, 40) B27 supplementation also exerted distinct effects on RPE metabolism that were dependent on the base medium. Antioxidants in B27, including selenium, vitamin E (tocopherol), catalase, and superoxide dismutase, can reduce oxidative stress and preserve mitochondrial function, supporting efficient ATP production through oxidative phosphorylation, particularly in low glucose conditions. While supplementation of galactose provides an alternative energy source, it may force cells to rely on oxidative phosphorylation as ATP is consumed in its initial metabolism, augmented further under conditions of low glucose.(41)

Several limitations should be acknowledged. For data normalization, cell culture surface area, protein concentration, and nuclei count are all commonly used approaches, yet each can yield distinct results. Because no consensus exists on whether cell number, size, or total protein content provides the most appropriate denominator to these data, there is no obvious solution other than to encourage the use of a standard normalization metric for the field to improve inter-study comparisons. While we report fractional changes to metabolites in culture medium to determine utilization, this form of quantitation places metabolites on the same scale, it can over-represent changes in metabolites that are in low abundance. For example, if the medium began with 200 μM of the molecule, a 10 μM reduction in ‘spent’ medium will reflect 5% consumption. But if the medium began with 10 μM of the molecule, 100% would be consumed. In both cases the actual consumption rate is the same. Metabolite consumption and production rates were approximated using two time points; however, metabolite consumption and production rates may not be linear within the sampling window, and earlier timepoints could reveal additional kinetic differences. Finally, medium composition is unlikely to be the only major determinant of RPE phenotype. Genetic factors, iPSC differentiation protocol variability, passage number, cell type (iPSC RPE vs fRPE), and a myriad other factors are also likely important determinants.

In static cell environments, nutrient composition is inherently dynamic, and no single medium formulation can maintain physiological concentrations of all metabolites throughout the culture period. Even within a given formulation, the metabolic state of RPE cells fluctuates over time,(14, 42) underscoring the lack of an ideal medium capable of continuously mimicking in vivo substrate supply. Rather than defining an optimal condition, this study provides a comparative framework detailing how existing media differentially influence RPE morphology, function, and metabolism. These findings emphasize the critical importance of media composition as an experimental variable and highlight the need to account for the nutrient microenvironment when interpreting RPE studies relevant to visual function and disease pathogenesis.

## MATERIALS AND METHODS

### Generation of patient-derived induced pluripotent stem cells (iPSCs)

Informed consent was obtained from all subjects prior to inclusion in the study (University of Washington IRB-approved STUDY00022435). Whole blood from normal control subjects was collected in Vacutainer CPT tubes (Becton Dickinson, Franklin Lakes, NJ, USA) for isolation of peripheral blood mononuclear cells (PBMC), which were then reprogrammed into iPSC using the CytoTune-iPS Sendai Reprogramming System (Thermo Fisher Scientific, Waltham, MA, USA).(*23, 24*) At least 2 iPSC clones were isolated from each patient, and all iPSC lines selected for study were of normal karyotype.

### Retinal pigmented epithelial (RPE) cell differentiation and human fetal RPE (fRPE) culture

iPSC cells were maintained in either mTeSR (StemCell Technologies, Vancouver, BC, Canada) or Essential 8^TM^ medium (Gibco, Thermo Fisher Scientific) and accutase passaged three times on Matrigel coated dishes before differentiation.(*23, 24*) At the end of the differentiation protocol, RPE cells identifiable by their characteristic cobblestone morphology and pigmentation were manually picked and expanded on Matrigel coated dishes. One to three RPE lines were generated from each of the patient-derived iPSC clone. RPE were cultured in MEMα containing FBS (Roche, Indianapolis, IN, USA), N1 supplement (1X, MilliporeSigma, Burlington, MA, USA), NEAA (1X, Gibco), hydrocortisone (20μg/l, MilliporeSigma), triiodo-thyronine (0.013μg/l, MilliporeSigma), taurine (250mg/l, MilliporeSigma) (THT) and penicillin-streptomycin (1X, Gibco). The first week of culture has 5% FBS and ROCKi (10μM, Selleck Chemicals, Houston, TX, USA), which was then reduced in 1% FBS without ROCKi.

To isolate fetal RPE, human fetal globes were obtained from the Birth Defects Research Laboratory (BDRL) at the University of Washington with ethics board approval and material written consent. This study was performed in accordance with ethical and legal guidelines of the University of Washington institutional review board. Globes were hemisected and the posterior eyecup revealed. The neural retina was removed, and RPE monolayer was subjected to 2% dispase digestion for 10 minutes at room temperature. RPE sheets were then carefully separated from underlying choroidal material, divided into smaller pieces and plated in MEMα based media as described above.(*43*) After 3 weeks, fRPE were trypsin passaged and expanded for experiments. fRPE from two donors (10 and 20 weeks) were used in this study.

All experimental cultures follow similar conditions. RPE were passaged using 0.25% trypsin, counted in the LUNA-II (Logos Biosystems, Annandale, VA, USA) and seeded in various media at 1.5×10^5^ cells/cm^2^ on Matrigel coated surfaces. The six media groups were kept on the same plate, when possible, to minimize plate to plate variation. Cells were maintained on 12 well plates in 1 ml media (Greiner Bio-One, Monroe, NC, USA), 8 well chambers in 300 μl media (Nunc, Thermo Fisher Scientific), 24 well PET Transwells in 200 μl apical and 900 μl basal media (Corning, Thermo Fisher Scientific) or 96 well XF Pro microplates in 80 μl media (Agilent Technologies, Santa Clara, CA, USA) in a 37°C incubator with 5% CO_2_, with media changes every other day (approximately 48hrs). RPE were used between passages 4 to 6, with at least n=3 per media group, and collected for assays after four to eight weeks in culture.

### Media formulation

Six different media used in this study (**Fig. 1A**) were formulated after literature review of RPE publications (**Supp. Table 1**). Medium 1 is the commonly used MEMα (Gibco #12561) based media described above, with 5% FBS during the first week, and 1% FBS during maintenance. Medium 2 is MEMα base with 2% B27 (Gibco #12587), a supplement favored as a serum-free alternative, along with NEAA and THT. Media 3 and 4 employs Human Plasma-Like Media (HPLM, Gibco #A4899101) designed to mimic the metabolic profile of human plasma.(*16*) Supplementation in media 3 and 4 is identical to that of media 1 and 2. Medium 5 is a high glucose formulation made of DMEM-HG (Gibco #10566) and Ham’s F-12 (Gibco #11765) in 70:30 ratio, with NEAA and 2% B27. FBS is given only in the first week – 10% in the first 2 days, 2% in the next five days. Medium 6 is X-VIVO-10 (Lonza #04-380Q, Basel, Switzerland), a serum-free cell medium for hematopoietic cells, with proprietary formulation. The only known components were obtained via phone conversation with Lonza – glucose (25mM) and glutamine (2mM). Complete media formulations with molarity of reported components are detailed in **Supp. Table 2**.

### Transepithelial electrical resistance (TER)

TER was measured using EVOM2 (World Precision Instruments, Sarasota, FL, USA) after leaving cells on bench for 10 min to equilibrate to room temperature. Two readings were taken from each well, and adjusted TER (Ω·cm^2^) was calculated by first subtracting values of empty filters then multiplying by surface area of transwells (0.33 cm^2^).

### Western blot and ELISA

RPE were cultured in 12 well plates for protein assessment. Conditioned media were collected 48 hours after media change for ELISA, and cells were scraped in MPER with Halt protease inhibitor (Thermo Fisher Scientific) for Western blot. To assess polarity of RPE on Transwells, apical and basal media were collected 24 hours after media change. VEGF, CFH (R&D Systems, Minneapolis, MN, USA) and ApoE (Abcam, Cambridge, MA, USA) ELISA were performed according to manufacturer’s instructions. For VEGF ELISA, apical and basal media were diluted 10- and 4-fold respectively. Conditioned media from 12 well dishes was diluted 5-fold for VEGF ELISA, 4-fold for CFH ELISA. Western blot was performed as previously described.(*23, 24*) Antibody dilutions are listed in **Supp. Table 3**.

### Immunostaining

RPE in 8 well chambers were fixed either with 4% PFA or chilled methanol for 10 min, and chambers in the former were permeabilized with 0.1% Triton X-100. Blocking was done with 5% donkey serum, and antibodies were diluted in 1% BSA. Primary antibodies (**Supp. Table 3**) were incubated overnight at 4°C. Cells were counterstained with DAPI (MilliporeSigma) and mounted with Fluoromount-G (SouthernBiotech, Birmingham, AL, USA). Imaging was performed on the Leica DM6000 (Leica Microsystems Inc., Deerfield, IL, USA). For REShAPE analysis, six images at 20X magnification were taken per well. Images with ZO-1 and F-actin only staining were exported as 16-bit greyscale TIF files for RPE segmentation.(*44*) To identify lipid droplets, live cell staining was performed where Lipi-Deep Red (Lipi-DR) with BODIPY-493/503 and Hoechst were co-incubated in iPSC RPE for 2h at 37°C. Six images at 63X magnification were automatically selected per well using a predefined acquisition pattern. In FIJI (RRID:SCR_002285) software (ImageJ, National Institutes of Health, Bethesda, MD, USA), BODIPY and Lipi-DR images were adjusted to remove background, converted to 16-bit, then auto thresholding performed with RenyiEntropy white method, followed by analyze particles, and summary data with %Area were plotted.

### Transmission electron microscopy

RPE on Transwells were processed for electron microscopy as described.(*23, 24*) Images taken at 20,000X on the JEM 1200EXII transmission electron microscope (JEOL USA, Inc., Peabody, MA, USA) were manually montaged into a panoramic view of the RPE monolayer. Three sections were imaged per filter, and sub-RPE deposits were quantified by tracing in FIJI software. Every image was analyzed with converted pixel to micrometer to obtain the exact area of deposits. Deposits were identified and individually outlined using the polygon tool by a masked observer.

### Mitochondria and glycolysis stress test

On the day of assay, two wells were stained with Hoechst-33258 (Thermo Fisher Scientific) and two representative areas per well were imaged on the EVOS microscope (Thermo Fisher Scientific) for nuclei counting. Subsequently, the same wells were collected in MPER lysis buffer for protein estimation. Mitochondria stress test and glycolysis stress test (n=5-6 per media group) were performed on the Seahorse XF Pro Analyzer (Agilent) in the University of Washington Diabetes Research Center - Metabolic and Cellular Phenotyping Core (NIDDK P30 DK017047). In brief, RPE were washed with XF DMEM + L-glutamine (2 mM) +/- glucose (5 mM) three times and placed in a CO_2_ null incubator for 1 hour. To assess mitochondria respiration, oligomycin (2 μM), FCCP (1 μM) and rotenone/antimycin A (2 μM) were added sequentially, with six readings taken between each drug. To assess glycolysis, RPE were incubated in media without glucose in the CO_2_ null incubator, followed sequential addition of glucose (5 mM), oligomycin (2 μM), and 2-deoxyglucose (50 mM) in the Seahorse Analyzer. Nuclei count from Hoechst images were extrapolated to the entire well with 0.114 cm^2^ area, and protein estimation was performed using Bradford assay and averaged for each well.

### Liquid chromatography with tandem mass spectrometry (LC-MS/MS)

Target metabolomics were performed on media and cell lysate from the same wells. Media were collected 24 and 48 hours after media change, and RPE were scraped into chilled 80% methanol as described.(*20, 21*) Blank media were run in parallel to obtain nutrient profile at 0 hours. LC-MS/MS was performed using a Shimadzu LC Nexera X2 UHPLC paired alongside a Sciex triple quadrupole QTRAP 5500 MS, as described previously.(*45*) Each metabolite was identified using mass and retention time listed in **Supp. Table 4**. To quantify metabolite consumption or production in the media, intensity values (obtained from the Sciex Multiquant 3.0.3 software) for the blank media were subtracted from the spent media values. To visualize intracellular metabolic changes on a heatmap, intensity values of each metabolite were normalized to that of media 1, then sorted left to right from highest and lowest fold change in each nutrient category.

## Statistical analysis

To compare amongst the six media groups, one-way ANOVA followed by Tukey’s post-hoc test was performed on GraphPad Prism 10 (RRID:SCR_002798) (GraphPad Software, Inc., La Jolla, CA, USA). When needed, pairwise comparison was done using T-test. A p-value of ≤ 0.05 was considered statistically significant. All data are presented as mean ± SEM. Targeted metabolomics data were analyzed using multivariate analysis by MetaboAnalyst 6.0 (RRID:SCR_015539) to obtain principal component distribution.

## Acknowledgments

The authors thank the Tom and Sue Ellison Stem Cell Core for their assistance in generating iPSC clones, Ed Parker for his electron microscopy expertise, and the Birth Defects Research Laboratory (funding provided by NIH under NICHD Grant # R24HD000836).

## Funding

National Institutes of Health Grant EY034364 (JRC and JD)

National Institutes of Health Grant EY034591(JRC and JD)

National Institutes of Health Grant EY032462 (JD)

National Institutes of Health Grant EY001730 NEI Vision Research Core (JBH, JRC)

Retina Research Foundation (JD)

BrightFocus Foundation M2020217 (JRC)

RPB Sybil B. Harrington Physician-Scientist Award for Macular Degeneration (JRC)

Unrestricted grant from Research to Prevent Blindness (JRC)

## Author contributions

Conceptualization: R.R.L., J.B.H., J.D., J.R.C.

Investigation: R.R.L., E.Z., Y.K., M.E., D.O., S.N., A.T., B.R.N., M.N., V.A.

Resources: D.O., K.B.

Formal analysis: R.R.L., D.T.H., J.B.H., J.D., J.R.C.

Supervision: J.B.H., J.D., J.R.C.

Writing—original draft: R.R.L., E.Z., D.T.H., J.R.C.

Writing—review & editing: R.R.L, D.T.H., J.B.H, K.B., J.D., J.R.C.

## Competing interests

The authors declare they have no competing interests.

## Data availability

All data needed to evaluate and reproduce the results in the paper are present in the paper and/or the Supplementary Materials. This study did not generate any new materials.

## Notes

### Competing Interest Statement

The authors have declared no competing interest.

### Summary of Updates

Title change, minor changes to text and citation numbers.

