## Supplementary Figures and Tables for "Nutrient Microenvironments Reprogram Pigment Epithelium Metabolism and Phenotype"

Supplementary Materials for

### Nutrient Microenvironments Determine Retinal Pigment Epithelium Metabolism and Phenotype

Rayne R. Lim *et al.*

**This PDF file includes:**

Figs. S1 to S7
Tables S1 to S4

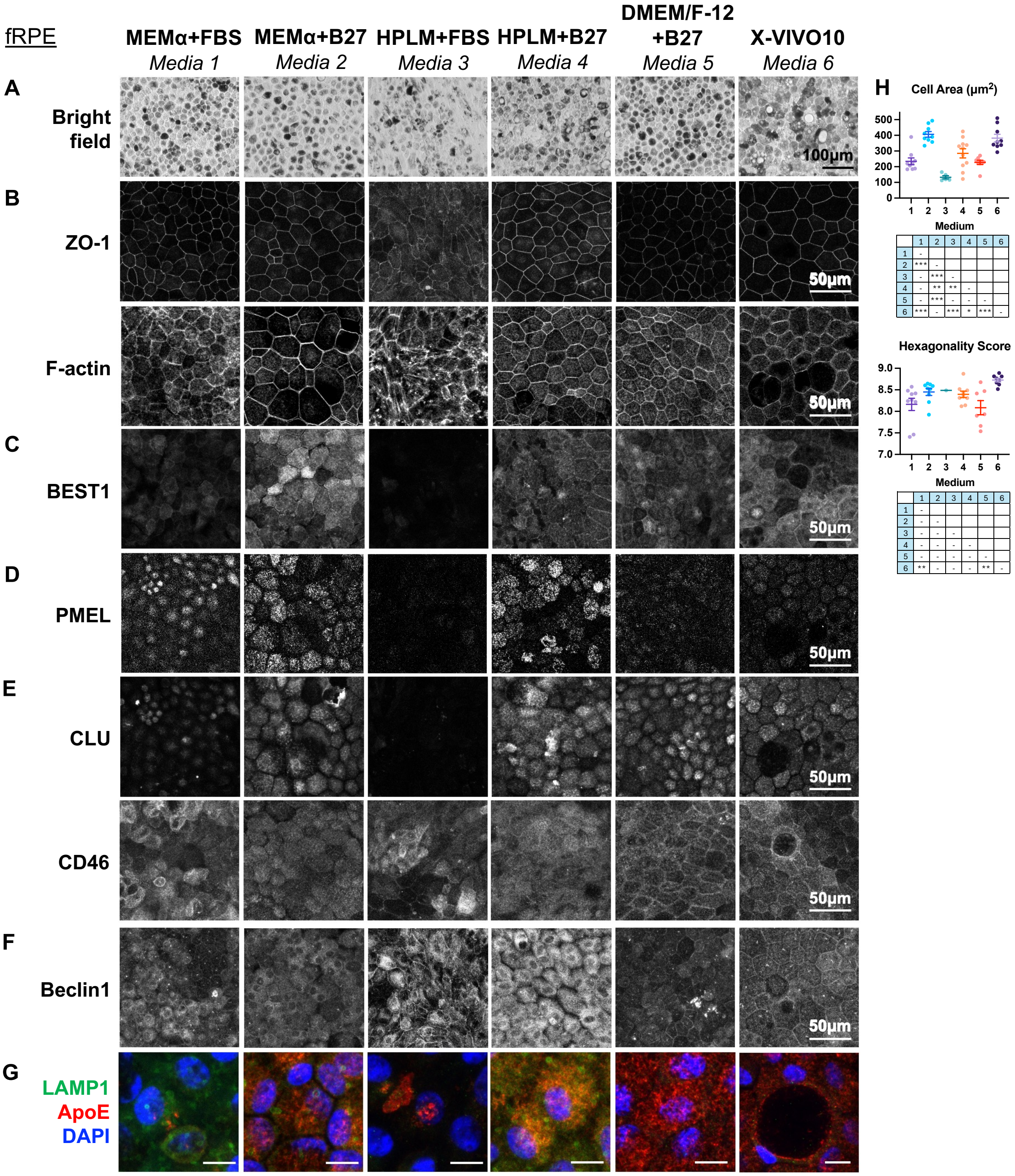


**Fig. S1. Brightfield and immunostaining images of fRPE cultured in Media 1 – 6.**

**(A)** Representative brightfield images of fRPE cultured on Matrigel-coated flat bottom dishes for 8 weeks. Immunostaining of fRPE on chamber slides for (**B**) ZO-1, F-actin (**C**) BEST1, (**D**) PMEL, (**E**) CLU, CD46, and (**F**) Beclin1. (**G**) Double staining of LAMP1and ApoE. Scale bar, 10μm. (**H**) fRPE cell area and hexagonality analyzed by REShAPE. Mean ± SEM. *, $p<0.05$; **, $p<0.01$; ***, $p<0.001$.

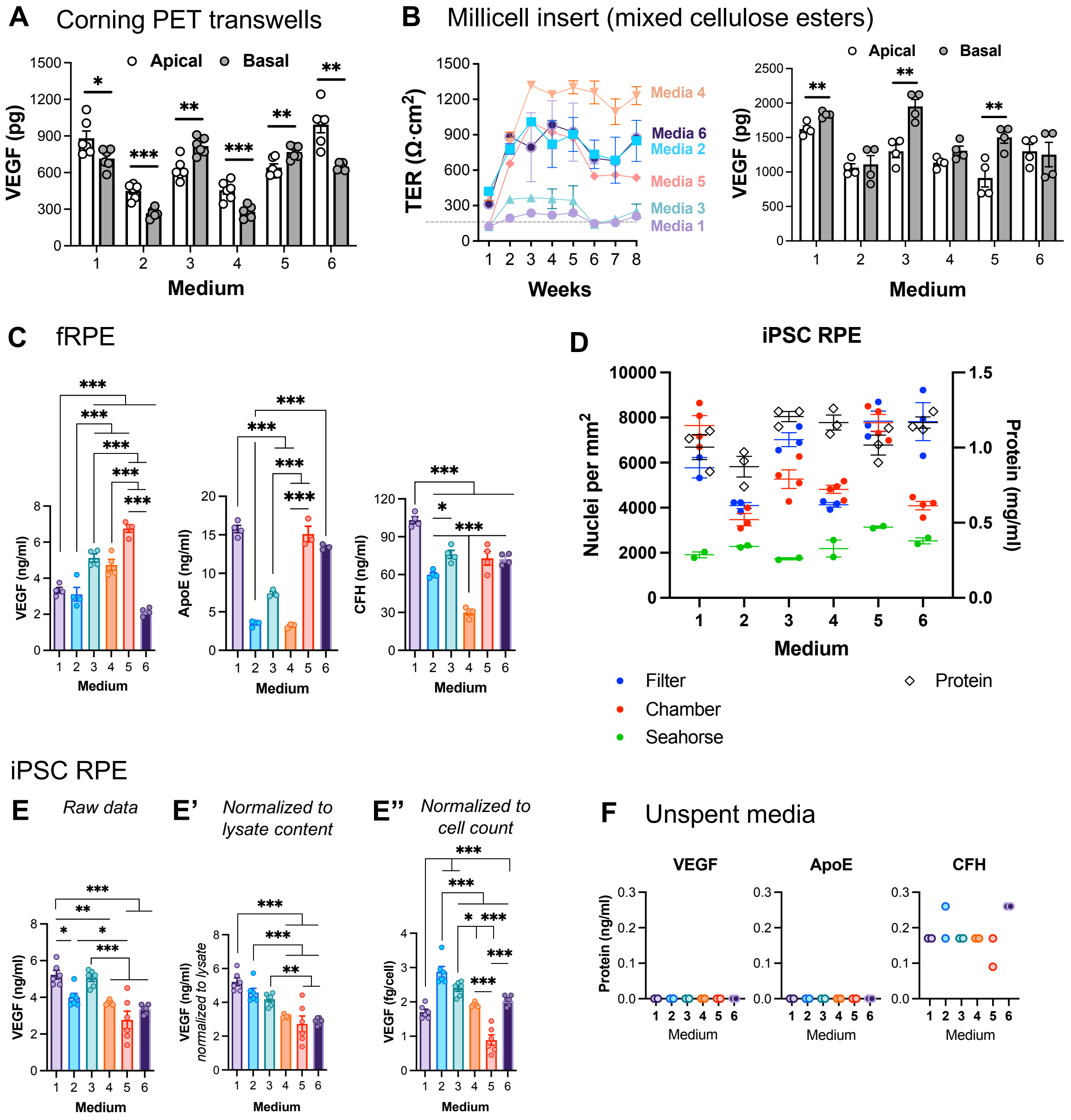


**Fig. S2. Polarized VEGF secretion, fRPE cytokine expression and normalization to nuclei count and protein content.** (**A**) VEGF ELISA of iPSC RPE cultured in PET Transwell filters at 8 weeks. (**B**) iPSC RPE cultured on mixed cellulose esters Millicell insert, and apical and basal VEGF secretion at 8 weeks. (**C**) fRPE cytokine expression on Matrigel-coated flat bottom dishes after 8 weeks of culture. (**D**) Hoechst-stained nuclei count of iPSC RPE cultured on PET filter, 8-well chamber and XFe96well plate, plotted with protein content from a 12-well dish. (**E**) iPSC RPE VEGF expression from Figure 2B normalized to lysate content or nuclei count. (**F**) VEGF, ApoE and CFH levels in unspent media.

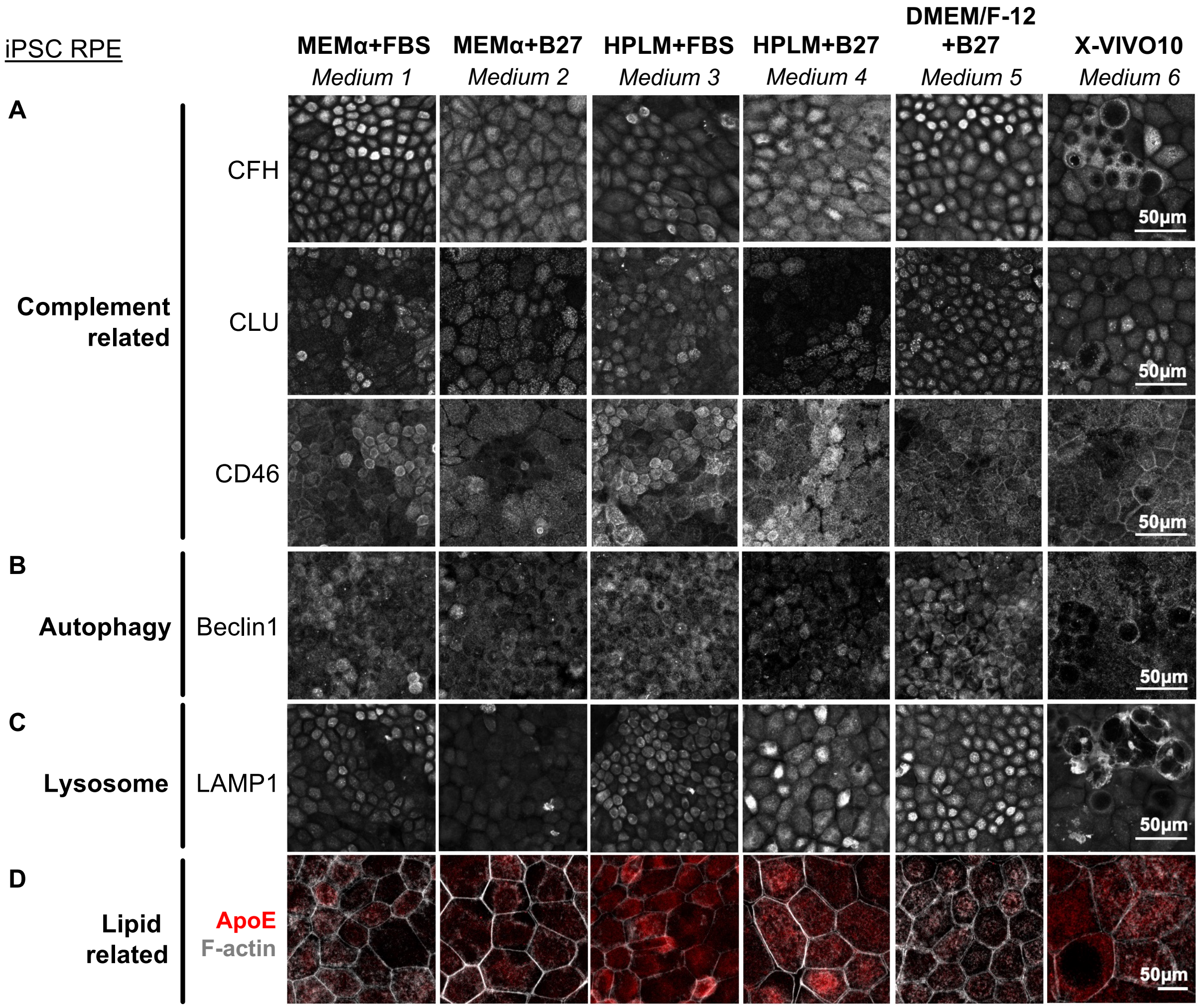


**Fig. S3. Immunostaining images of iPSC RPE cultured in Media 1 – 6.** Immunostaining of 8-week-old iPSC RPE on chamber slides for (**A**) complement related CFH, CLU, CD46, (**B**) autophagy related Beclin1, (**C**) Lysosomal LAMP1, and (**D**) double staining of ApoE with F-actin.

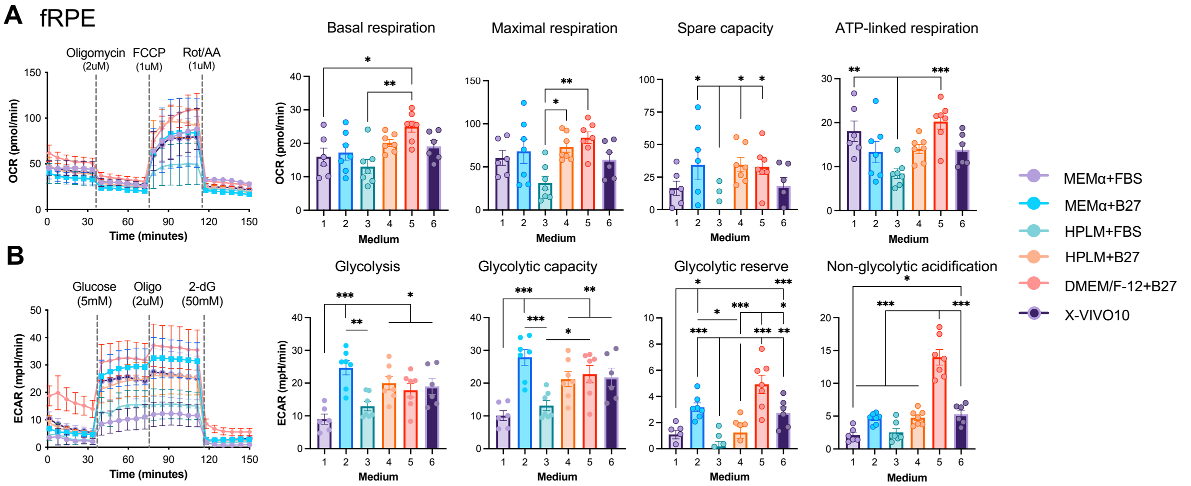


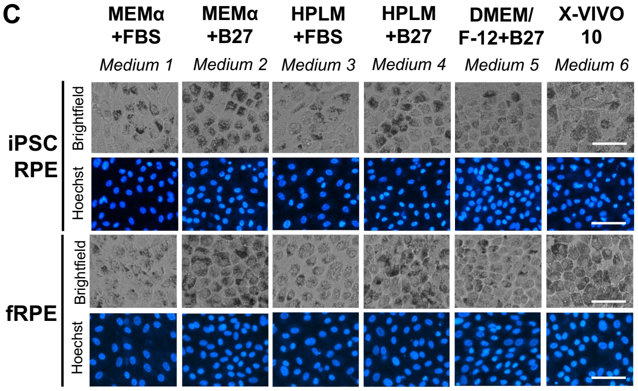


D

| Media | iPSC RPE Protein (mg/ml) | iPSC RPE Cell count (per mm²) | fRPE Protein (mg/ml) | fRPE Cell count (per mm²) |
|---|---|---|---|---|
| 1 | 0.42 | 1910 | 0.39 | 1239 |
| 2 | 0.17 | 2283 | 0.23 | 2589 |
| 3 | 0.36 | 1736 | 0.42 | 1723 |
| 4 | 0.14 | 2187 | 0.28 | 2154 |
| 5 | 0.32 | 3140 | 0.39 | 2519 |
| 6 | 0.33 | 2527 | 0.40 | 2245 |

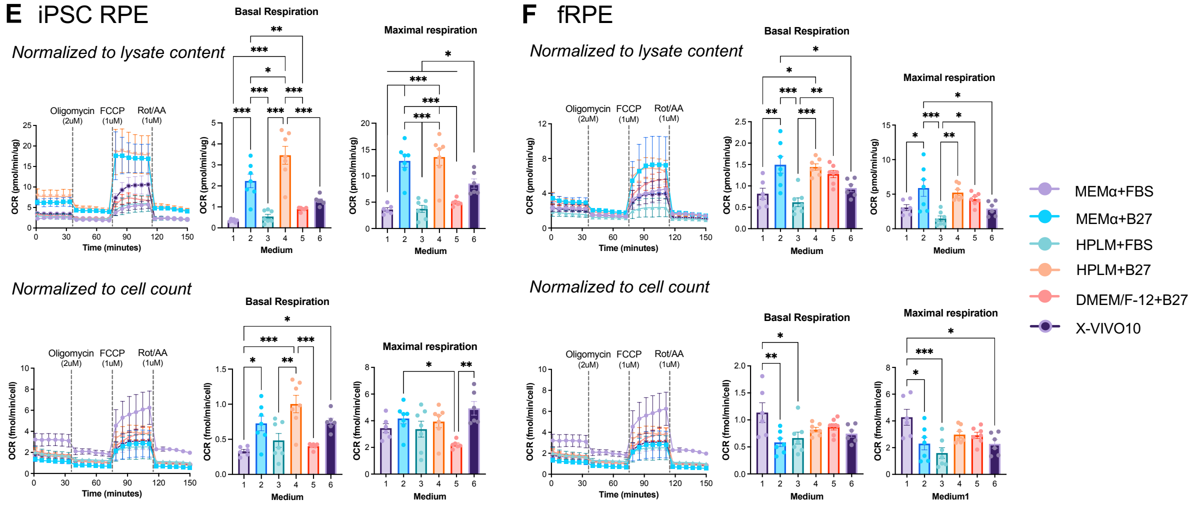


**Fig. S4. OCR and ECAR data normalized to lysate content and cell count.** (**A,B**) fRPE were seeded and maintained on Matrigel coated XFe96 well plates in respective Media 1 – 6 for four weeks prior to mitochondrial and glycolysis stress test. (**C**) Brightfield images and Hoechst-stained nuclei of iPSC RPE and fRPE cultured on Matrigel coated XFe96 well after 4 weeks. Scale bar, 50µm. (**D**) Protein concentration and cell count per well of RPE maintained in Media 1 – 6 on XFe96 well plate. (**E,F**) iPSC RPE and fRPE oxygen consumption rate (OCR) data normalized to lysate content and cell count. Mean ± SEM. *, $p<0.05$; **, $p<0.01$; ***, $p<0.001$.

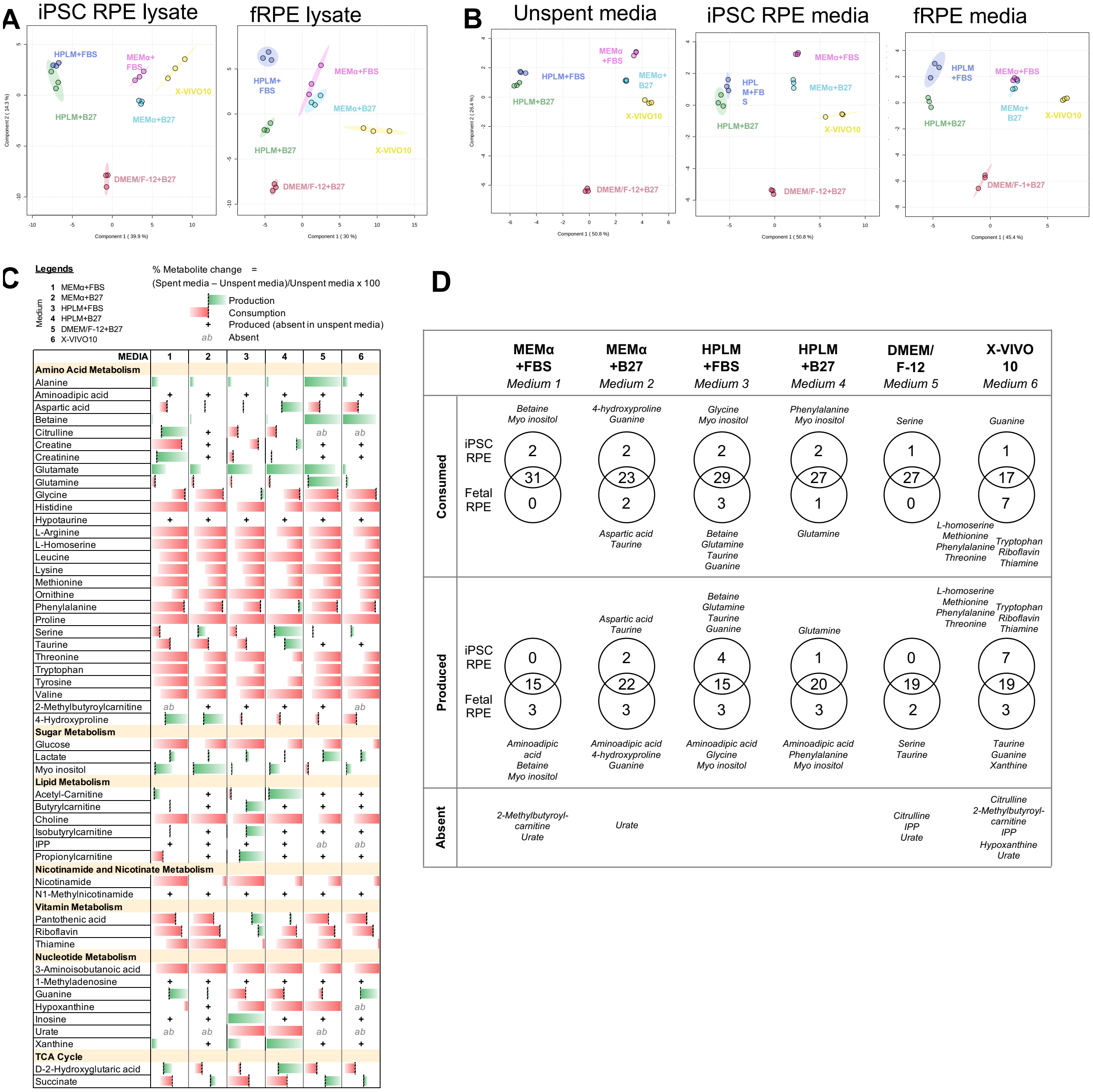


**Fig. S5. PLS-DA, fRPE extracellular metabolite, consumption and production patterns across Media 1 – 6.** Multi-variate analysis with Partial Least Squares Discriminant Analysis (PLS-DA) of (**A**) RPE lysate after 8 weeks culture, as well as (**B**) unspent and 48h spent media in both iPSC RPE and fRPE. (**C**) Metabolite changes in fRPE media analyzed as described in Figure 4. (**D**) Venn diagram depicting commonly and uniquely consumed or produced metabolites in iPSC RPE and fRPE within each media type.

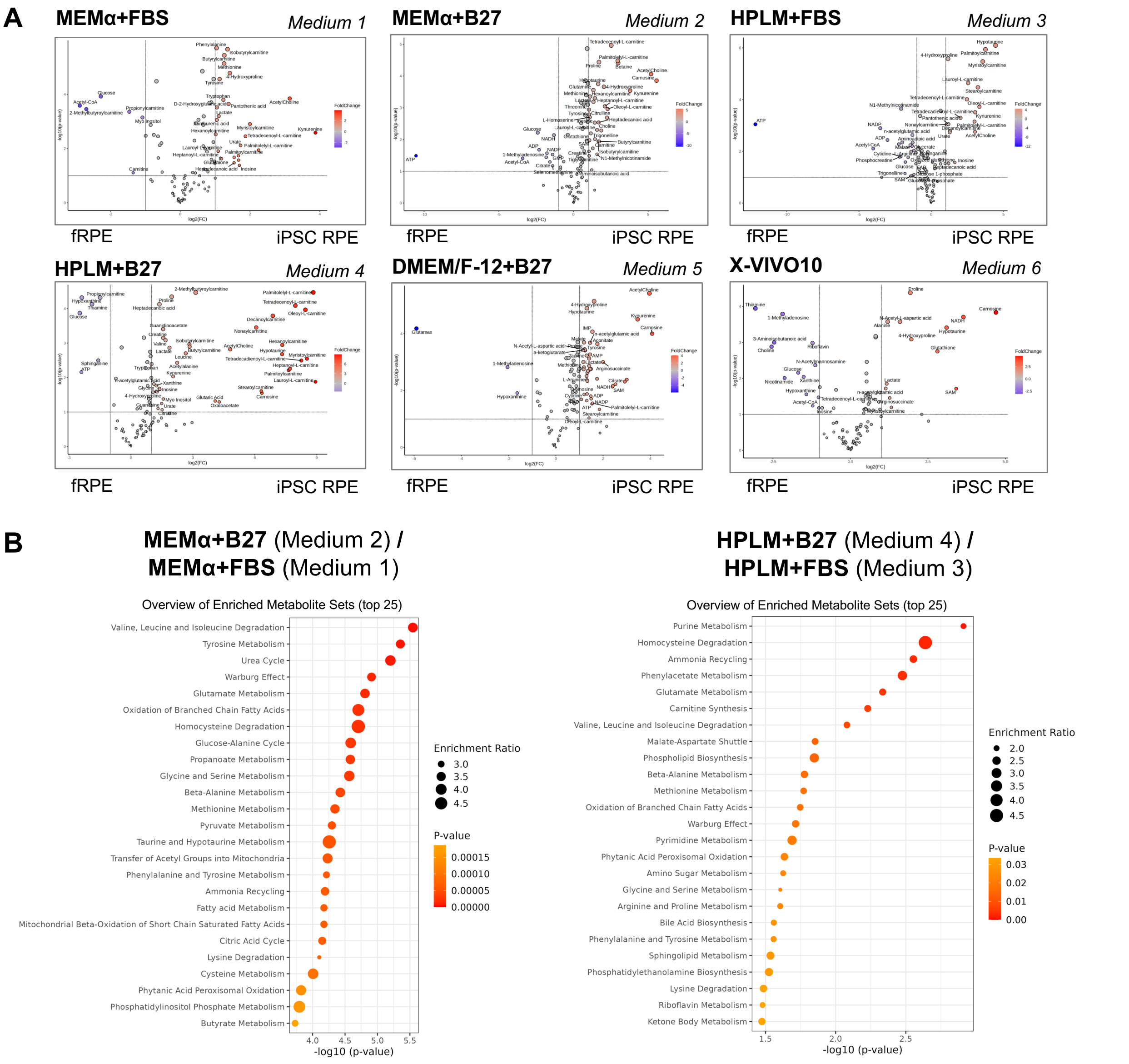


**Fig. S6. Volcano plots of intracellular metabolite and enriched metabolite sets.** (**A**) Volcano plots comparing intracellular metabolites between fRPE and iPSC RPE within each media type. (**B**) Enrichment analysis in MetaboAnalyst 6.0 showed overview of top 25 enriched metabolite sets in Medium 1 vs 2 and Medium 3 vs 4.

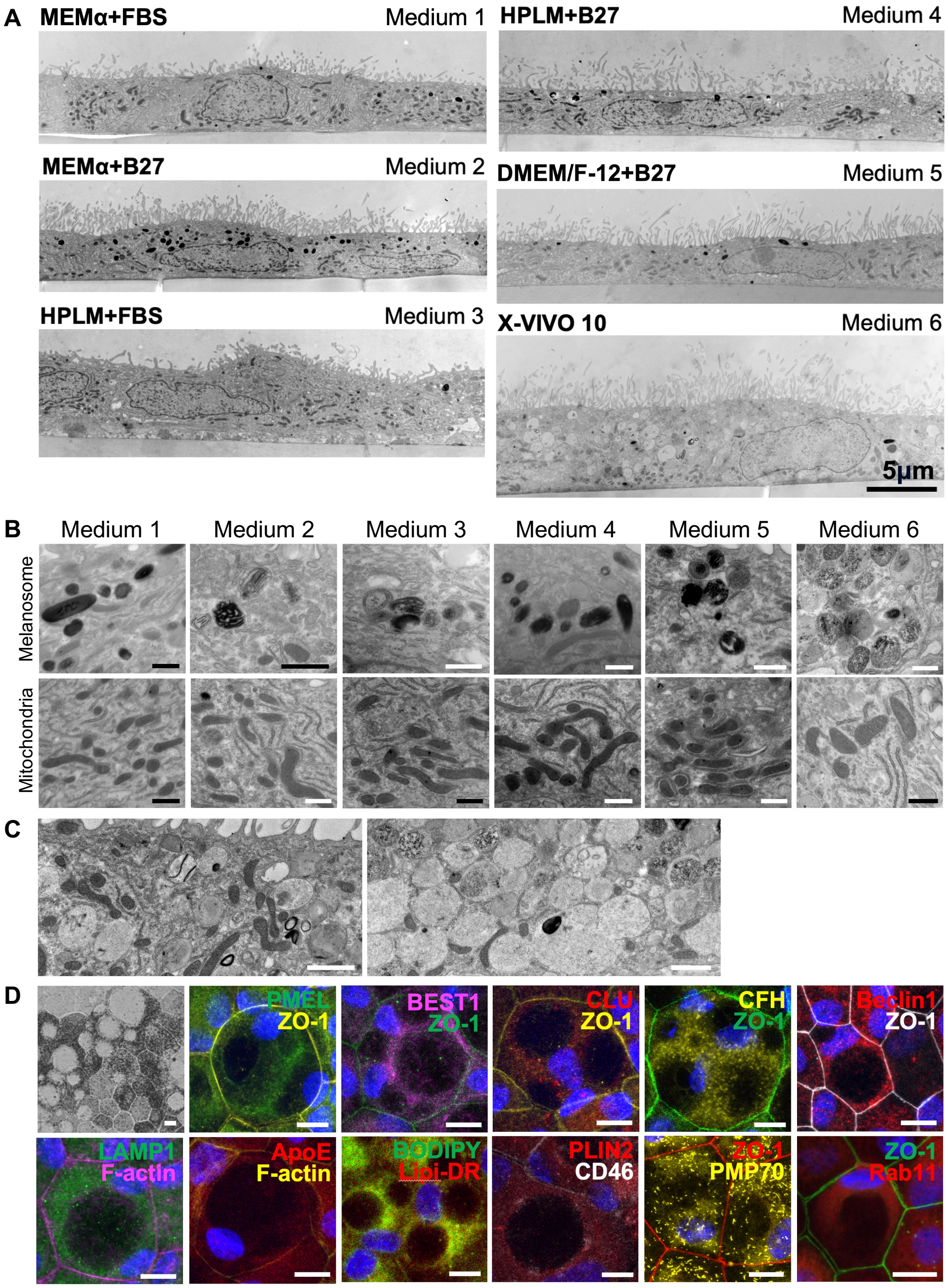


**Fig. S7. TEM of iPSC RPE in Media 1 – 6, immunostaining of vacuoles in iPSC RPE cultured in Media 6.** (**A**) TEM panoramic images of iPSC RPE cultured for 8-weeks on PET Transwell filters. (**B**) Representative high magnification TEM images of melanosomes and mitochondria in RPE maintained in Media 1 – 6. Scale bar, 500nm. (**C**) iPSC RPE cultured in Media 6 developed numerous vacuoles that were not electron lucent, with some containing remnants of cytosolic organelles resembling autophagosomes. Scale bar, 1μm. (**D**) Vacuoles were visible under bright field imaging, and immunostaining with various markers did not uncover identity of vacuolar contents. Scale bar, 10μm.

| Media Type | Glucose (mM) | RPE cell type | Differentiation method | RPE Matrix | Media supplements | | | | | Culture (weeks) | Seeding density (cells/$cm^2$) | Filter type | TEER ($\Omega \cdot cm^2$) | Subtracted empty transwell? | PMID |
|---|---|---|---|---|---|---|---|---|---|---|---|---|---|---|---|
| | | | | | NEAA | N1/N2 | THT | FBS | Others | | | | | | |
| MEMα | 5.5 | fRPE | - | Human ECM | ✓ | ✓ | ✓ | 5% | | NR | $1.78 \times 10^{5}$ | PET | 500 | - | 16877436 |
| | | fRPE | - | Fibronectin | ✓ | ✓ | ✓ | 1% | Glutamine (2mM) | 4 | $0.89 \times 10^{5}$ | PET | 400-600 | - | 26990160 |
| | | iPSC-RPE | Directed | Matrigel | ✓ | ✓ | ✓ | 1% | | 6 | $2.0 \times 10^{5}$ | PET | 200 | Yes | 34929159 |
| | | iPSC-RPE | Directed | - | ✓ | ✓ | ✓ | 5% | | 6 | - | PET | 170-200 | - | 24873859 |
| | | iPSC-RPE | Directed | Human ECM | ✓ | ✓ | ✓ | 5% | | 15 | - | PET | 400-800 | Yes | 27400791 |
| | | iPSC-RPE | Directed | Vitronectin | ✓ | ✓ | ✓ | 5% | PGE2 (50uM) | NR | $2.2 \times 10^{5}$ | - | - | - | 35880133 |
| | | iPSC-RPE | Directed | Vitronectin | ✓ | ✓ | ✓ | 5% | Glutamine (2mM)<br>PGE2 (50uM) | 8 | - | PET | 260 | - | 30651323 |
| | | iPSC-RPE | Directed | Vitronectin | ✓ | ✓ | ✓ | 5% | Glutamine (2mM) | 6 | $2.2 \times 10^{5}$ | PET | 200-1500 | - | 34887495 |
| | | iPSC-RPE | Directed | Vitronectin | ✓ | | ✓ | 5% | | 6 | - | PET | 400-1000 | - | 34911940 |
| | | iPSC-RPE | Spontaneous | Laminin | ✓ | ✓ | ✓ | 1% | GlutaMAX (2mM)<br>normal human AB serum (4%)<br>Nicotinamide (10mM) | 6 to 8 | $1.66 \times 10^{5}$ | PET | 200 | Yes | 28969679 |
| DMEM-LG + Ham's F-12 (67:29) | ~6.7 | iPSC-RPE | Directed | CELLstart | | | | | B27 (1.9%)<br>L-glutamine (1.9mM) | 7 | $5 \times 10^{5}$ | PET | 600 | Yes | 31376722 |
| DMEM/F-12 + MEMα (1:1) | ~11.5 | iPSC-RPE | Directed | Human placental ECM | ✓ | ✓ | ✓ | 2% | Glutamine (1mM)<br>Sodium Pyruvate (1mM) | 4 | $0.89 \times 10^{5}$ | PET | 200 | - | 28132833 |
| DMEM/F-12 | 17.5 | iPSC-RPE | Directed | Matrigel | ✓ | | | | KOSR (4%)<br>β-mercaptoethanol (50μM) | 6 | $4.5 \times 10^{5}$ | PET | 200-300 | Yes | 37331647 |
| | | iPSC-RPE | Directed | Synthemax II | ✓ | ✓ | | 10% | GlutaMAX (2mM)<br>Sodium pyruvate (1mM)<br>Nicotinamide (10mM) | 4 | $0.89 \times 10^{5}$ | PET | 250-300 | - | 36092714 |
| DMEM-HG + Ham's F-12 (3:1) | ~20.5 | iPSC-RPE | Spontaneous | Laminin | ✓ | | | | B27, no retinoic acid (2%) | - | $0.66 \times 10^{5}$ | PET | 350-450 | Yes | 24030465 |
| | | iPSC-RPE | Spontaneous | Laminin | ✓ | | | | B27, no retinoic acid (2%) | 12 | $1.66 \times 10^{5}$ | PET | 200 | Yes | 28878022 |
| | | iPSC-RPE | Spontaneous | Laminin | ✓ | | | | B27, no retinoic acid (2%) | 4 | - | PET | 200 | Yes | 31840941 |
| DMEM-HG | 25 | Porcine RPE | - | Laminin/ Entactin | | | | 1% | Glutamine (100mM) | 6 | $3.0 \times 10^{5}$ | PET | 700-800 | Yes | 28687758 |
| | | Porcine RPE | - | Collagen IV | ✓ | | | 1% | | 3 | $3.0 \times 10^{5}$ | PET | 400 | Yes | 24861273 |
| | | iPSC-RPE | Directed | Matrigel | | | | 1% | GlutaMAX<br>Sodium pyruvate | 4 | $1.0 \times 10^{5}$ | PET | - | - | 30572641 |
| X-VIVO 10 | 25 | iPSC-RPE | Directed | Matrigel | | | | | | 8 | $6.0 \times 10^{5}$ | PET | 200 | - | 31890732 |
| | | iPSC-RPE | Directed | Matrigel | | | | | | 5 | $1.33 \times 10^{5}$ | PET | - | - | 33918210 |
| | | hESC-RPE | Directed | Matrigel | | | | | | 4 | - | PET | - | - | 25604686 |
| Knockout DMEM | 25 | iPSC-RPE | Spontaneous | Matrigel | ✓ | | | | KOSR (15%)<br>GlutaMAX (2mM)<br>β-mercaptoethanol (0.1mM) | 7 to 9 | $2.1 \times 10^{5}$ | PET | 100-300 | - | 34202702 |
| | | iPSC-RPE | Spontaneous | Matrigel | ✓ | | | | KOSR (15%)<br>GlutaMAX (2mM)<br>β-mercaptoethanol (0.1mM) | 8 to 10 | $2.1 \times 10^{5}$ | PET | 500-750 | Yes | 37137441 |

**Table S1. Summary of published RPE culture, method of differentiation and reported TER.**

| B27 | Base media | MEMα | MEMα | HPLM | HPLM | DMEM/F-12 (3:1) | X-VIVO 10 |
|---|---|---|---|---|---|---|---|
| | NEAA | ✓ | ✓ | ✓ | ✓ | ✓ | |
| | N1 | ✓ | | ✓ | | | |
| | THT | ✓ | ✓ | ✓ | ✓ | | |
| | B27 | | 2% | | 2% | 2% | |
| | FBS (*not included in table*) | 1% | | 1% | | | |
| | Pen/Strep | ✓ | ✓ | ✓ | ✓ | ✓ | ✓ |
| **B27** | **MEDIUM** | **1** | **2** | **3** | **4** | **5** | **6** |
| | **Proteinogenic Amino Acids (mM)** | | | | | | |
| | L-Alanine | 0.38 | 0.38 | 0.53 | 0.53 | 0.13 | - |
| | L-Arginine | 0.60 | 0.60 | 0.11 | 0.11 | 0.58 | - |
| | L-Asparagine | 0.44 | 0.44 | 0.15 | 0.15 | 0.13 | - |
| | L-Aspartic acid | 0.33 | 0.33 | 0.12 | 0.12 | 0.13 | - |
| | L-Cysteine | 0.57 | 0.57 | 0.040 | 0.040 | 0.20 | - |
| | L-Glutamate | NA | NA | 0.08 | NA | NA | - |
| | L-Glutamic acid | 0.61 | 0.61 | 0.18 | 0.18 | 0.13 | |
| | L-Glutamine | 2.00 | 2.00 | 0.55 | 0.55 | 3.10 | 4.00 |
| | Glycine | 0.78 | 0.78 | 0.40 | 0.40 | 0.41 | - |
| | L-Histidine | 0.20 | 0.20 | 0.11 | 0.11 | 0.17 | - |
| | L-Isoleucine | 0.40 | 0.40 | 0.070 | 0.070 | 0.57 | - |
| | L-Leucine | 0.40 | 0.40 | 0.16 | 0.16 | 0.59 | - |
| | L-Lysine | 0.40 | 0.40 | 0.20 | 0.20 | 0.62 | - |
| | L-Methionine | 0.10 | 0.10 | 0.030 | 0.030 | 0.15 | - |
| | L-Phenylalanine | 0.19 | 0.19 | 0.080 | 0.080 | 0.29 | - |
| | L-Proline | 0.46 | 0.46 | 0.30 | 0.30 | 0.19 | - |
| | L-Serine | 0.35 | 0.35 | 0.25 | 0.25 | 0.41 | - |
| | L-Threonine | 0.40 | 0.40 | 0.14 | 0.14 | 0.59 | - |
| | L-Tryptophan | 0.049 | 0.049 | 0.060 | 0.060 | 0.058 | - |
| | L-Tyrosine | 0.20 | 0.20 | 0.080 | 0.080 | 0.29 | - |
| | L-Valine | 0.39 | 0.39 | 0.22 | 0.22 | 0.59 | - |
| | **Non-proteinogenic Amino Acids (mM)** | | | | | | |
| | Alpha-Aminobutyrate | NA | NA | 0.020 | 0.020 | NA | - |
| | L-Citrulline | NA | NA | 0.040 | 0.040 | NA | - |
| | L-Cystine | 0.10 | 0.10 | 0.10 | 0.10 | NA | - |
| | 4-Hydroxy-L-proline | NA | NA | 0.020 | 0.020 | NA | - |
| | L-Ornitine | NA | NA | 0.070 | 0.070 | NA | - |
| | **Amino Acids Derivatives (mM)** | | | | | | |
| | L-Acetyl glycine | NA | NA | 0.090 | 0.090 | NA | - |
| ✓ | Glutathione (reduced) | NA | 0.0033 | 0.025 | 0.028 | 0.0033 | - |
| | Taurine | 2.00 | 2.00 | 2.09 | 2.09 | NA | - |
| | N-Trimethylglycine (betaine) | NA | NA | 0.070 | 0.070 | NA | - |
| | **Vitamins (mM)** | | | | | | |
| | p-Aminobenzoate | NA | NA | 0.0070 | 0.0070 | NA | - |
| ✓ | DL Alpha Tocopherol Acetate | NA | 0.0021 | NA | 0.0021 | 0.0021 | - |
| ✓ | DL Alpha-Tocopherol (Vit E) | NA | 0.0024 | NA | 0.0024 | 0.0024 | - |
| | Ascorbic Acid | 0.28 | 0.28 | NA | NA | NA | - |
| ✓ | D-Biotin | 0.00041 | 0.011 | 0.00082 | 0.011 | 0.010 | - |
| | Choline | 0.0071 | 0.0071 | 0.022 | 0.022 | 0.050 | - |
| | Folic Acid | 0.0023 | 0.0023 | 0.0023 | 0.0023 | 0.0072 | - |
| | myo-Inositol/i-inositol | 0.011 | 0.011 | 0.19 | 0.19 | 0.060 | - |
| | Niacinamide | 0.0082 | 0.0082 | 0.0082 | 0.0082 | 0.023 | - |
| | D-Calcium pantothenate | 0.0021 | 0.0021 | 0.00052 | 0.00052 | 0.0062 | - |
| | Pyridoxine | 0.0049 | 0.0049 | 0.0049 | 0.0049 | 0.014 | - |
| | Riboflavin | 0.00027 | 0.00027 | 0.0010 | 0.0010 | 0.00080 | - |
| | Thiamine | 0.00297 | 0.00297 | 0.0030 | 0.0030 | 0.0086 | - |
| | Vitamin B12 | 0.0010 | 0.0010 | 0.0000037 | 0.0000037 | 0.00030 | - |

**Legend**

| Concentration (mM) | |
|---|---|
| High | |
| Medium | |
| Low | Absent |

| B27 | Base media | MEMα | MEMα | HPLM | HPLM | DMEM/F-12 (3:1) | X-VIVO 10 |
|---|---|---|---|---|---|---|---|
| | NEAA | ✓ | ✓ | ✓ | ✓ | ✓ | |
| | N1 | ✓ | | ✓ | | | |
| | THT | ✓ | ✓ | ✓ | ✓ | | |
| | B27 | | 2% | | 2% | 2% | |
| | FBS (*not included in table*) | 1% | | 1% | | | |
| | Pen/Strep | ✓ | ✓ | ✓ | ✓ | ✓ | ✓ |
| **B27** | **MEDIUM** | **1** | **2** | **3** | **4** | **5** | **6** |
| | **Other Components (mM)** | | | | | | |
| | Acetate | NA | NA | 0.040 | 0.040 | NA | - |
| | Acetone | NA | NA | 0.060 | 0.060 | NA | - |
| ✓ | BSA, fatty acid free Fraction V | NA | 0.038 | NA | 0.038 | 0.038 | - |
| ✓ | Carnitine (L-carnitine in B27) | NA | 0.012 | 0.040 | 0.052 | 0.012 | - |
| ✓ | Catalase | NA | 0.000 | NA | 0.000 | 0.000 | - |
| | Citrate | NA | NA | 0.130 | 0.130 | NA | - |
| ✓ | Corticosterone | NA | 0.058 | NA | 0.058 | 0.058 | - |
| | Creatine | NA | NA | 0.040 | 0.040 | NA | - |
| | Creatinine | NA | NA | 0.075 | 0.075 | NA | - |
| | D-Glucose | 5.56 | 5.56 | 5.00 | 5.00 | 20.50 | 25.00 |
| ✓ | Ethanolamine | NA | 0.00081 | NA | 0.00081 | 0.00081 | - |
| | Formate | NA | NA | 0.050 | 0.050 | NA | - |
| | Fructose | NA | NA | 0.040 | 0.040 | NA | - |
| ✓ | Galactose | NA | 0.08 | 0.06 | 0.14 | 0.08 | - |
| | Glycerol | NA | NA | 0.12 | 0.12 | NA | - |
| ✓ | Human transferrin | 0.000063 | 0.000063 | 0.000063 | 0.000063 | 0.000063 | - |
| | Hydrocortisone | 0.000055 | 0.000055 | 0.000055 | 0.000055 | NA | |
| | 2-Hydroxybutyrate | NA | NA | 0.050 | 0.050 | NA | - |
| | 3-Hydroxybutyrate | NA | NA | 0.050 | 0.050 | NA | - |
| | Hypoxanthine | NA | NA | 0.010 | 0.010 | 0.010 | - |
| | Lactate | NA | NA | 1.60 | 1.60 | NA | - |
| ✓ | Linoleic acid | NA | 0.0036 | NA | 0.0036 | 0.0037 | - |
| ✓ | Linolenic acid | NA | 0.0036 | NA | 0.0036 | 0.0036 | - |
| | Lipoic Acid | 0.0010 | 0.0010 | NA | NA | 0.0003 | - |
| | Malonate | NA | NA | 0.01 | 0.01 | NA | - |
| | Phenol Red | 0.027 | 0.027 | 0.014 | 0.014 | 0.029 | - |
| ✓ | Progesterone | 0.000023 | 0.000020 | 0.000023 | 0.000020 | 0.000020 | - |
| ✓ | Putrescine 2HCl | 0.182 | 0.183 | 0.182 | 0.183 | 0.183 | - |
| | Pyruvate | NA | NA | 0.050 | 0.050 | NA | - |
| ✓ | Recombinant Human Insulin | 0.0009 | 0.0005 | 0.0009 | 0.0005 | 0.0005 | - |
| | Sodium Pyruvate | 1.00 | 1.00 | NA | NA | 0.30 | - |
| | Succinate | NA | NA | 0.020 | 0.020 | NA | - |
| ✓ | Superoxide Dismutase (KU/ml) | NA | 0.38 | NA | 0.38 | 0.38 | - |
| ✓ | Triiodo-Thyronin (T3) | 2.00E-08 | 0.0031 | 2.00E-08 | 0.0031 | 0.0031 | - |
| | Urate | NA | NA | 0.35 | 0.35 | NA | - |
| | Urea | NA | NA | 5.00 | 5.00 | NA | - |
| | **Inorganic Salts (mM)** | | | | | | |
| | Ammonium Chloride | NA | NA | 0.04 | 0.04 | NA | - |
| | Calcium Chloride | 1.80 | 1.80 | 2.35 | 2.35 | 1.35 | - |
| | Calcium Nitrate | NA | NA | 0.04 | 0.04 | NA | - |
| | Magnesium Chloride | NA | NA | 0.48 | 0.48 | 0.18 | - |
| | Magnesium Sulfate | 0.81 | 0.81 | 0.35 | 0.35 | 0.57 | - |
| | Potassium Chloride | 5.33 | 5.33 | 4.10 | 4.10 | 4.63 | - |
| | Potassium Phosphate Monobasic | NA | NA | 0.015 | 0.015 | NA | - |
| | Sodium Bicarbonate | 26.19 | 26.19 | 24.00 | 24.00 | 35.03 | - |
| | Sodium Chloride | 117.24 | 117.24 | 105.00 | 105.00 | 116.55 | - |
| | Sodium Phosphate Monobasic | 1.01 | 1.01 | 0.87 | 0.87 | 0.63 | - |
| | Sodium Phosphate dibasic (NA2HPO4) anhydrous | NA | NA | NA | NA | 0.30 | - |
| | Thymidine | NA | NA | NA | NA | 0.00087 | - |
| | **Trace Elements (mM)** | | | | | | |
| | Cupric Sulfate | NA | NA | NA | NA | 0.000003 | - |
| | Ferric Nitrate | NA | NA | NA | NA | 0.000173 | - |
| | Ferric Sulfate | NA | NA | NA | NA | 0.00090 | - |
| ✓ | Sodium Selenite | 0.000029 | 0.000072 | 0.000029 | 0.000072 | 0.000072 | - |
| | Zinc Sulfate | NA | NA | NA | NA | 0.00090 | - |

**Table S2. Complete media formulations of reported components in molarity.**

| Target | Company | Catalogue # | ICC dilution | WB dilution | Cell fixation |
|---|---|---|---|---|---|
| ApoE | MilliporeSigma | AB947 | 1:1000 | 1:1500 | 4% PFA |
| β-actin | GeneTex | GTX629630 | - | 1:5000 | - |
| Beclin1 | Proteintech | 66665-1-Ig | 1:100 | - | Methanol |
| BEST1 | MilliporeSigma | MAB5466 | 1:50 | - | 4% PFA |
| | Novus Biologicals | NB300-164 | - | 1:100 | - |
| BODIPY 493/503 | MilliporeSigma | 790389 | 50μg/ml | - | Live cell imaging |
| CD46 | BioRad | MCA2113 | 1:200 | - | 4% PFA |
| CFH | Quidel | A229 | 1:50 | - | 4% PFA |
| CLU | MilliporeSigma | AB825 | 1:200 | - | 4% PFA |
| CRALBP | gift from Dr Jing (19) | | - | 1:500 | - |
| EZR | Cell Signaling Technologies | 3145 | - | 1:1000 | - |
| F-actin-568 | Invitrogen | A12380 | 1:200 | - | 4% PFA |
| LAMP1 | Proteintech | 21997-1-AP | 1:200 | - | 4% PFA |
| Lipi-Deep Red | Dojindo | LD04 | 0.2μM | - | Live cell imaging |
| LRAT | Abcam | ab137304 | - | 1:500 | - |
| PLIN2 | Proteintech | 15294-1-AP | 1:200 | - | 4% PFA |
| PMEL | Invitrogen | MA5-13232 | 1:50 | 1:50 | 4% PFA |
| PMP70 | MilliporeSigma | SAB4200181 | 1:200 | - | 4% PFA |
| Rab11 | BD Laboratories | 610656 | 1:200 | - | 4% PFA |
| RPE65 | MilliporeSigma | MAB5428 | - | 1:2500 | - |
| TYR | Abcam | ab170905 | - | 1:1000 | - |
| ZO-1 | Invitrogen | 40-2200 | 1:100 | - | 4% PFA |

**Table S3. Antibodies used in immunocytochemistry and Western blot.**

| Metabolite | CAS ID | Q1 Mass (Da) | Q3 Mass (Da) | Retention time (RT) |
|---|---|---|---|---|
| 1-Methyladenosine | 15763-06-1 | 282.1 | 150.1 | 1.85 |
| N1-Methylnicotinamide | 3106-60-3 | 138 | 78 | 1.63 |
| 2-Methylbutyroylcarnitine | 31023-25-3 | 246.2 | 85.1 | 1.3 |
| 3-Aminoisobutanoic acid | 144-90-1 | 104.1 | 58 | 1.37 |
| 4-Hydroxyproline | 51-35-4 | 132.1 | 86 | 2.63 |
| N-Alpha-acetyllysine | 152473-69-3 | 189.1 | 84.1 | 5 |
| N-Acetylglutamic acid | 1188-37-0 | 190.1 | 84.1 | 3.02 |
| cis-Aconitic acid | 499-12-7 | 173 | 85 | 3.66 |
| Adenine | 73-24-5 | 134 | 107 | 0.73 |
| Adenosine | 58-61-7 | 268.1 | 136 | 0.79 |
| Adipic acid | 124-04-9 | 145 | 83 | 3.14 |
| ADP | 58-64-0 | 426 | 79.1 | 4.6 |
| AICAR | 3031-94-5 | 259.1 | 111 | 0.99 |
| a-ketoglutarate | 328-50-7 | 145 | 101 | 3.02 |
| L-Alanine | 56-41-7 | 90 | 44 | 2.35 |
| Aminoadipic acid | 542-32-5 | 160.1 | 116.1 | 3.75 |
| Adenosine monophosphate | 61-19-8 | 346 | 134 | 3.76 |
| Argininosuccinic acid | 2387-71-5 | 291.1 | 69.9 | 4.6 |
| L-Aspartic acid | 56-84-8 | 134.1 | 74 | 3.49 |
| Adenosine triphosphate | 987-65-5 | 507.9 | 136.1 | 5.69 |
| Betaine | 590-46-5 | 118.1 | 58 | 1.67 |
| Butyrylcarnitine | 25576-40-3 | 232.2 | 85.1 | 1.5 |
| Carnosine | 305-84-0 | 227.1 | 110.1 | 3.61 |
| Choline | 62-49-7 | 104.1 | 60.1 | 1.51 |
| Citraconic acid | 498-23-7 | 129 | 85 | 3.63 |
| Citric acid | 77-92-9 | 191.1 | 87 | 2.39 |
| Citrulline | 372-75-8 | 174.1 | 131.1 | 3.28 |
| Creatine | 57-00-1 | 132.1 | 90 | 2.66 |
| Creatinine | 60-27-5 | 114 | 44 | 0.8 |
| L-Cystathionine | 56-88-2 | 221.1 | 79.1 | 0.57 |
| L-Cystine | 56-89-3 | 239 | 74 | 4.26 |
| Cytidine | 65-46-3 | 244 | 112.1 | 1.39 |
| D-2-Hydroxyglutaric acid | 103404-90-6 | 149.1 | 77 | 1.37 |
| Decanoyl-L-carnitine | 3992-45-8 | 316.3 | 85.1 | 0.6 |
| FAD | 146-14-5 | 784.1 | 437.1 | 3.8 |
| D-Fructose | 57-48-7 | 179.1 | 71 | 1.24 |
| Glucose 1-phosphate | 59-56-3 | 259 | 79 | 4.17 |
| Glucose 6-phosphate | 56- 73-5 | 259 | 79 | 4.11 |
| Gamma-Aminobutyric acid | 56-12-2 | 104.1 | 87 | 2.57 |
| D-Glucose | 492-62-6 | 179 | 89 | 1.95, 2.07 |

| | | | | |
|---|---|---|---|---|
| L-Glutamic acid | 56-86-0 | 148.1 | 84.1 | 3.3 |
| L-Glutamine | 56-85-9 | 147.1 | 84.1 | 3.03 |
| Glutaric acid | 110-94-1 | 131 | 87 | 3.41 |
| Glycine | 56-40-6 | 76 | 30.2 | 2.57 |
| Guanosine monophosphate | 85-32-5 | 362.1 | 79 | 4.22 |
| Glutathione | 70-18-8 | 306 | 143.1 | 3.8 |
| Oxidized Glutathione | 13081-14-6 | 611 | 306 | 5.56 |
| Guanosine triphosphate | 86-01-1 | 521.9 | 159 | ~5.2 |
| Guanine | 73-40-5 | 152 | 110 | 1.21 |
| Guanosine | 118-00-3 | 282.1 | 150 | 1.64 |
| Heptadecanoic acid | 506-12-7 | 269.101 | 135.1 | 1.05 |
| Hexanoylcarnitine | 22671-29-0 | 260.2 | 85.1 | 0.9 |
| L-Histidine | 332-80-9 | 156.1 | 110 | 3.35 |
| Hypotaurine | 300-84-5 | 108.1 | 64 | 2.44 |
| Hypoxanthine | 68-94-0 | 135 | 65 | 0.8 |
| Inosinic acid | 131-99-7 | 347 | 79 | 4 |
| Inosine | 58-63-9 | 267 | 135 | 1.1 |
| Isopentenyl pyrophosphate | 18687-43-9 | 245.1 | 79 | 2.61 |
| Isobutyryl-L-carnitine | 25518-49-4 | 232.1 | 85.1 | 1.5 |
| L-Kynurenine | 343-65-7 | 207.1 | 144 | 1.59 |
| L-Acetylcarnitine | 14992-62-2 | 204.1 | 85 | ~1.86 |
| Lactate | 50-21-5 | 89 | 43 | 1.13 |
| L-Arginine | 74-79-3 | 175.1 | 70.1 | 4.28 |
| L-Asparagine | 70-47-3 | 133.1 | 70.1 | 2.94 |
| L-Leucine | 61-90-5 | 132.1 | 86 | 1.65 |
| L-Homoserine | 1927-25-9 | 120.1 | 74 | 3.04 |
| Linoleic acid | 60-33-3 | 281.1 | 221 | 0.5 |
| L-Lysine | 56-87-1 | 147.1 | 84 | 4.23 |
| L-Malic acid | 6915-15-7 | 133 | 115 | 3.55 |
| L-Methionine | 63-68-3 | 150.1 | 61 | 1.65 |
| myo-Inositol | 87-89-8 | 179 | 87 | 2.9 |
| Myristoyl-L-carnitine | 25597-07-3 | 372.4 | 85.1 | 0.4 |
| N1-Methylnicotinamide | 3106-60-3 | 137 | 78 | 1.85 |
| N-Acetyl-L-aspartic acid | 997-55-7 | 176.1 | 74 | 3.12 |
| NAD | 53-84-9 | 664 | 136 | 4.33 |
| NADP | 53-59-8 | 744 | 136 | 5.36 |
| N-alpha-Acetyl-L-lysine | 152473-69-3 | 187.1 | 145.1 | 3.48 |
| Niacinamide | 98-92-0 | 123 | 80 | 0.41 |
| Nicotinamide D4 | 347841-88-7 | 127 | 84 | 0.45 |
| Nicotinamide riboside | 2181-04-6 | 256.1 | 123 | 2.25 |
| Ornithine | 70-26-8 | 133 | 70 | 4.3 |

| | | | | |
|---|---|---|---|---|
| Oleoyl-L-carnitine | 38677-66-6 | 426.4 | 85.1 | 0.5 |
| L-Palmitoylcarnitine | 1985-18-8 | 400.4 | 85.1 | 0.4 |
| Pantothenic acid | 137-08-6 | 218.1 | 71 | 1.13 |
| L-Phenylalanine | 63-91-2 | 166.1 | 120 | 1.37 |
| Phosphoserine | 407-41-0 | 186 | 70 | 5 |
| Phosphocreatine | 19333-65-4 | 210 | 79 | 4.42 |
| L-Proline | 4298 08 2 | 116.1 | 70.1 | 2.1 |
| Propionylcarnitine | 17298-37-2 | 218.1 | 85.1 | 1.9 |
| Pyridoxal | 66-72-8 | 166 | 79 | 0.44 |
| Pyridoxamine | 85-87-0 | 169.1 | 134.1 | 1.61 |
| Pyroglutamic acid | 98-79-3 | 130.1 | 84 | 2.02 |
| Riboflavin | 83-88-5 | 377.1 | 243.1 | 1.07 |
| S-Adenosylmethionine | 86867-01-8 | 399.3 | 136 | 4.5 |
| C5H11NO2Se | 3211-76-5 | 196.99 | 194.6 | 97 |
| L-Serine | 56-45-1 | 106 | 60 | 2.83 |
| Stearoylcarnitine | 25597-09-5 | 428.5 | 85.1 | 0 |
| Succinic acid | 110-15-6 | 117 | 73 | 2.92 |
| Taurine | 107-35-7 | 126 | 108 | 1.8 |
| Thiamine | 59-43-8 | 265 | 122.1 | 2.1 |
| L-Threonine | 72-19-5 | 120.1 | 102 | 2.5 |
| Trigonelline | 535-83-1 | 138 | 92 | 1.81 |
| Tiglylcarnitine | 64681-36-3 | 244.2 | 85.1 | 1.36 |
| L-Tryptophan | 73-22-3 | 205 | 146 | 1.79 |
| L-Tyrosine | 60-18-4 | 182.1 | 136 | 1.92 |
| Uridine diphosphate glucose | 133-89-1 | 611 | 499 | 4.59 |
| Uric acid | 69-93-2 | 167 | 124 | 2.6 |
| L-Valine | 72-18-4 | 118.1 | 72 | 1.76 |
| Xanthine | 69-89-6 | 151 | 108 | 1.1 |
| udp glucosamine | 17479-04-8 | 606 | 385 | 4.28 |

**Table S4. LCMS mass and retention time.**
